# Sensitizing tumor response to topoisomerase I antibody drug conjugate by selective CDK7 inhibition

**DOI:** 10.1101/2025.11.23.690049

**Authors:** Hyerim Jung, Yoonji Lee, Donghoon Yu, Yeejin Jeon, Hwankyu Kang, Dahun Um, Mooyoung Seo, Dongsik Park, Jeongjun Kim, Seung-Joo Lee, Jaeseung Kim, Lauren Escobedo, Yilun Sun, Anish Thomas, Kiyean Nam, Tae-Kyung Kim

**Affiliations:** Department of Life Sciences, Pohang University of Science and Technology, Pohang, Gyeongbuk, 37673, Korea; Qurient Co., Ltd., C801, 242 Pangyo-ro, Bundang-gu, Seongnam-si, Gyeonggi-do, 13487, Korea; University of Maryland Marlene & Stewart Greenebaum Comprehensive Cancer Center, University of Maryland School of Medicine, Baltimore, MD 21201, USA; Developmental Therapeutics Branch, Center for Cancer Research, National Cancer Institute, NIH, Bethesda, MD 20892, USA; Institute for Convergence Research and Education in Advanced Technology, Yonsei University, Seoul, 03722, Republic of Korea

## Abstract

This study investigates the transcriptional impact of Q901, a highly selective CDK7 inhibitor in clinical development. Q901 primarily disrupted MYC and E2F-dependent transcription program, downregulating cell cycle control and DNA damage repair pathways. CDK7 binding at the promoter-proximal regions was dramatically stabilized by Q901, leading to reduced occupancy of MYC, E2F, and RNA Polymerase II (RNAPII). These findings offered a novel therapeutic strategy to enhance cancer susceptibility to TOP1-DNA protein crosslinks (TOP1-DPCs) induced by TOP1 inhibitors. Resistance to TOP1 inhibitors arises through activation of DNA repair pathway when elongating RNAPII encounters TOP1-DPCs. By suppressing RNAPII transition from initiation to elongation and DNA repair pathways, Q901 stabilizes TOP1-DPCs and sensitizes tumor to TOP1 inhibitors. Preclinical studies demonstrated enhanced tumor suppression when combining Q901 with TOP1 inhibitor-based antibody-drug conjugates (TOP1i-ADCs), highlighting its potential as a therapeutic option for cancers resistant to TOP1i-ADC therapy.

**Teaser:** Overcoming ADC cross-resistance through combination approach.

## Introduction

Cyclin-dependent kinase 7 (CDK7), as part of the CDK-activating kinase (CAK) complex (comprising CDK7, cyclin H, and MAT1), plays a pivotal role in regulating both cell cycle and transcription (*1, 2*). The CAK complex facilitates the phosphorylation of the T-loop in CDK1, CDK2, CDK4, and CDK6, which is essential for their activation leading to cell cycle progression (*3*). In transcription, CDK7 functions as a core component of the transcription factor II H (TFIIH) complex, where it plays critical roles in both the initiation and elongation phases of RNA polymerase II (RNAPII)-mediated transcription (*4–7*). During initiation, CDK7 phosphorylates the Ser5 residue of the RNAPII C-terminal domain (CTD), a process that facilitates promoter escape (*4, 5*). In the elongation phase, CDK7 phosphorylates CDK9, which in turn phosphorylates the pausing factors NELF and DSIF, as well as the RNAPII CTD at the Ser2 residue (*6*). This phosphorylation leads to the release of NELF, the conversion of DSIF into a positive elongation factor, and the continuation of transcription elongation (*7*). Additionally, these phosphorylation events are crucial for recruiting capping enzymes and splicing factors, which regulate RNA processing (*8*). Thus, CDK7-mediated phosphorylation of RNAPII is integral to the regulation of transcription and RNA processing. The dual role of CDK7 in cell cycle regulation and transcription underscores its importance as a fundamental regulator of cellular processes.

Cancer is characterized by complex molecular alterations, including changes in gene expression mediated by transcription factors, chromatin remodelers, or histone-modifying enzymes (*9*). Targeting each of these factors individually is challenging due to their complexity and sheer number. Instead, the general transcriptional machinery, which is essential for gene transcription, has emerged as a viable therapeutic target. CDK7, as a key component of this machinery, presents an attractive target for cancer therapy. CDK7 expression is often elevated in many types of cancer with poor prognosis (*10, 11*). Previously developed CDK7 inhibitors, such as THZ1 and SY-1365, have demonstrated anticancer effects by suppressing RNAPII CTD phosphorylation and inducing global transcriptional downregulation. However, their cytotoxic effects were attributed to off-target activity on CDK12 and CDK13 (*12, 13*). In contrast, a newer inhibitor YKL-5-124, which exhibits greater specificity for CDK7 without affecting other CDKs, did not significantly reduce RNAPII CTD phosphorylation or cause global transcriptional downregulation (*14*). THZ1 and its derivatives, SY-1365 and SY-5609, were shown to arrest the cell cycle at the G2/M phase and induce apoptosis (*12, 13, 15*), whereas YKL-5-124 primarily induced G1 cell cycle arrest without noticeable apoptosis (*14*). Similarly, genetic disruption of CDK7 did not result in global transcriptional downregulation or inhibition of CTD phosphorylation (*16*).

Topoisomerase I (TOP1) is a critical enzyme that alleviates torsional stress in DNA strands during replication and transcription by transiently introducing single-strand breaks into DNA. During this process, TOP1 forms a covalent bond between its catalytic tyrosine (Tyr) residue and the 3’ end of the broken DNA backbone, generating transient and reversible Topoisomerase I cleavage complexes (TOP1ccs). TOP1 inhibitors, such as topotecan (TPT) and irinotecan, stabilize TOP1ccs and convert them into irreversible TOP1 DNA-protein crosslinks (TOP1-DPCs), thereby blocking the progression of transcription and replication forks, ultimately leading to DNA damage and subsequent cell death (*17, 18*). TOP1 inhibitors are widely used as payloads in antibody-drug conjugates (ADCs) (*19*), a therapeutic approach that combines the specificity of monoclonal antibodies with the cytotoxic effects of small-molecule drugs (*20*). Trastuzumab deruxtecan (T-DXd), an-FDA-approved TOP1 inhibitor-based ADC (TOP1i-ADC) showed an impressive objective response rate (ORR) of 60.9% and median progression-free survival (PFS) of 16.4 months in patients with HER2-positive metastatic breast cancer who had been previously treated with multiple lines of HER2-targeted therapies (*21*). More recently, T-DXd has shown significant survival benefits in HER2-low metastatic breast cancer, expanding its use beyond HER2 positive patient population (*22*). To further enhance the efficacy of TOP1i-ADC conjugates, drug combination strategies are also being actively explored. A combination of TOP1i-ADC with either a DNA damage response inhibitor (e.g., PARP or ATR inhibitor) or immunotherapy (e.g., anti-PD-1 antibody) has been shown to increase the anticancer efficacy of TOP1i-ADC (*23, 24*).

Theoretically, cancer cells with high transcription and replication rates are expected to be more susceptible to TOP1 inhibitors. However, a potential limitation of TOP1 inhibitor therapy lies in the ability of cells to repair DNA damage through transcription-coupled repair (TCR). Studies have shown that collisions between elongating RNAPII and TOP1-DPCs can activate the ubiquitin-proteasome pathway, leading to the degradation of both complexes, followed by DNA repair (*18*). A recent study revealed that the sensitivity of rapidly replicating small cell lung cancer (SCLC) cells to TPT can be enhanced when combined with the CDK7 inhibitor, THZ1. Mechanistically, targeting transcription with THZ1 reduces collisions between RNAPII and TOP1-DPCs, thereby preventing the proteasomal degradation of TOP1-DPCs (*25*). This combination represents a novel strategy to increase the efficacy of TOP1 inhibitors, particularly in the context of balancing the therapeutic efficacy and toxicity of TOP1i-ADCs. However, a deeper understanding of the mechanism of action is needed to successfully translate the combination of CDK7 inhibitors and TOP1i-ADCs into clinical applications.

The present study explores the molecular mechanism of CDK7 inhibition in transcriptional regulation using Q901 (mocaciclib), a novel and highly selective CDK7 inhibitor. Q901 covalently binds to Cys312 within the ATP-binding pocket of CDK7, effectively suppressing its kinase activity without affecting other CDKs. By selectively inhibiting CDK7, Q901 traps CDK7 and disrupts the RNAPII transcription process at promoter-proximal regions of genes involved in MYC- and E2F-driven oncogenic transcription programs. Q901 prevents RNAPII from transitioning into stable elongation, leading to the stabilization of TOP1-DPCs, which provides a potential mechanism for overcoming cancer resistance to TOP1 inhibitors. Preclinical studies further demonstrated that Q901 synergizes with TOP1i-ADCs, resulting in enhanced tumor suppression. These findings underscore Q901’s potential as a promising therapeutic option, particularly for cancers resistant to TOP1i-ADC therapies.

## Results

### Q901 is a highly selective covalent inhibitor of CDK7

Q901 is a covalent CDK7 inhibitor featuring a pyrazolo[1,5-a][1,3,5]triazine as a hinge binder and an α-fluoro acrylamide as its covalent binding moiety (Fig. 1A and fig. S1A). It forms a covalent bond with cysteine 312 (Cys312) within the ATP binding pocket of CDK7 in a competitive manner. Mass spectrometry analysis demonstrated that the intensity of the cysteine-containing peptide RPNCPVETL (residues 309-317) was reduced by approximately 99% following Q901 treatment. In contrast, the intensity of another cysteine-containing peptide, RPGPTPGCQLP (residues 298-308), remained unaffected, indicating a site-specific interaction between Q901 and Cys312 (Fig. 1B and fig. S1B).

**Fig. 1.**
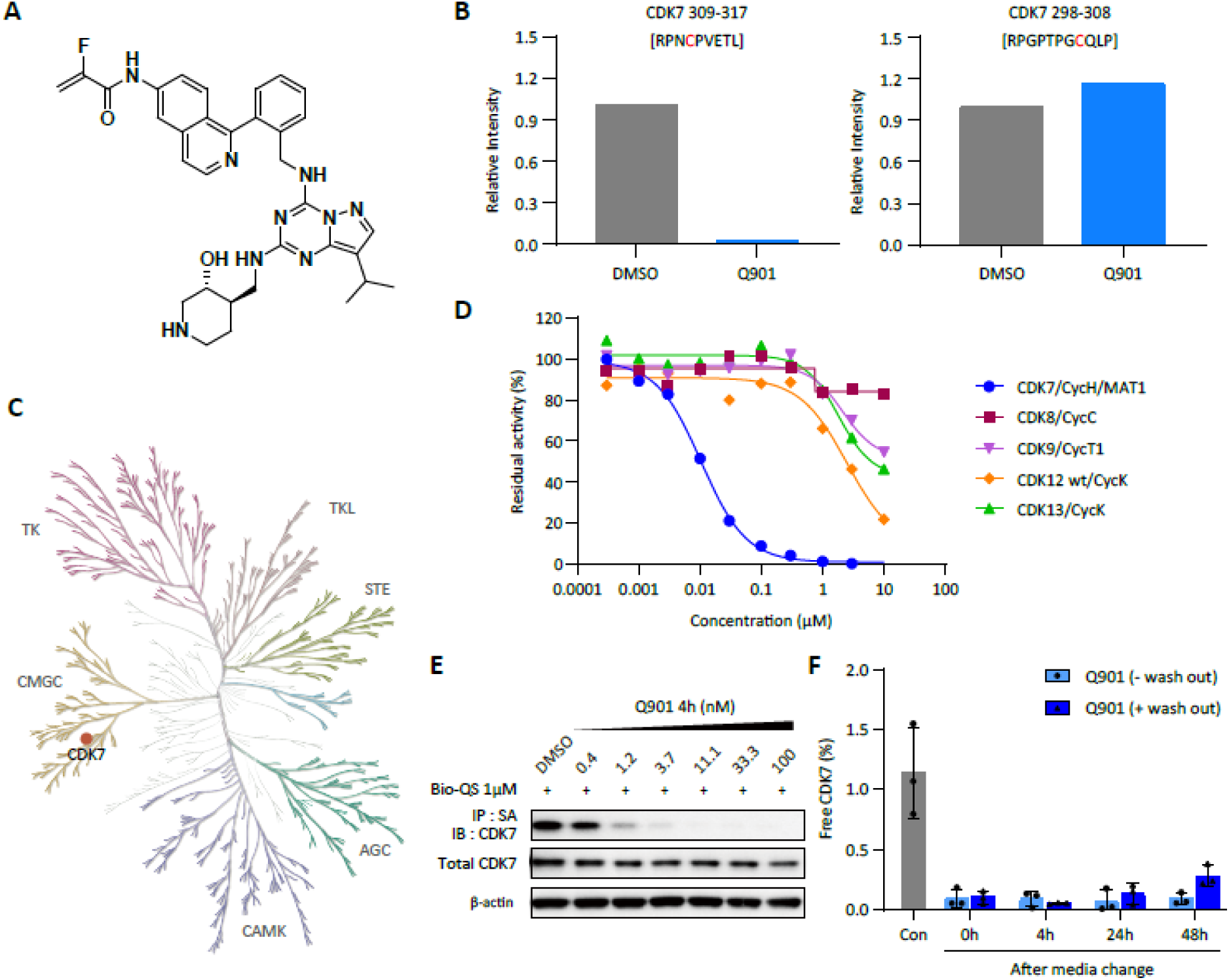
Q901 is a CDK7 selective covalent inhibitor. (**A**) Chemical structure of Q901. (**B**) Assessment of the covalent binding site was conducted via mass spectrometry analysis of the recombinant CDK7-cyclin H-MAT1 (CAK) trimetric complex. The recombinant CAK complex was incubated with Q901 or DMSO for 1 h, digested for 18 h, and analyzed by mass spectrometry. (**C**) KinMap image illustrating the kinase inhibition profile of Q901 against a panel of 410 kinases (397 protein kinase assays and 13 lipid kinase assays). The inhibition profile was determined by measuring the residual activity at 1 μM for 1 h using the PanQinase® Activity Assay. ATP concentration was set at the apparent ATP-Km value for each kinase. A red dot indicates 99% inhibition of CDK7 by Q901 at this concentration. (**D**) Efficacy and selectivity of Q901 against other CDKs at ATP concentrations corresponding to the apparent ATP-Km value for each kinase. Residual activity (%) was measured after a 1 h incubation with Q901 at the indicated concentrations. (**E**) CDK7 target occupancy assay using Bio-QS, a biotinylated analog of Q901. Cell lysates were prepared from A2780 cells treated with Q901 or DMSO for 4 h at the indicated concentrations and subjected to immunoprecipitation (IP) using Bio-QS and streptavidin agarose beads (SA). IP samples and whole-cell lysates were immunoblotted with an anti-CDK7 antibody. (**F**) Washout-based target occupancy assay to measure the duration of CDK7 inhibition. A2780 cells treated with 6 nM Q901 for 4 h were washed with fresh medium and incubated for the indicated times. Cells were lysed, treated with Bio-QS, and immunoprecipitated with streptavidin agarose beads (SA). The percentage of free CDK7 was calculated by normalizing CDK7 levels in IP samples from Q901 treatment to those in IP samples from the DMSO-treated group (n = 3; two-way ANOVA with Tukey’s multiple comparisons test, data represent mean ± SD).

*In vitro*, Q901 inhibited CDK7 activity with an IC_50_ of 10 nM but exhibited much lower inhibitory activity against other kinases in an inhibition profiling assay with a panel of 410 kinases, including thirteen lipid kinases (Fig. 1, C and D, and table S1). The inhibition by Q901 was highly selective for CDK7, demonstrating a residual activity value of 1 (99% inhibition) at a concentration of 1 μM. In comparison, the IC_50_ values for other CDKs were significantly higher than that for CDK7, ranging from 242-fold to over 1,000-fold (Table S1). Additionally, Q901 binding to CDK7 remained stable even after washout in a target occupancy assay, consistent with its covalent binding activity (Fig. 1, E and F, and fig. S1C).

Special attention was given to the selectivity of Q901 against CDK8, CDK9, CDK12, and CDK13, as these CDKs are also involved in transcriptional regulation. Inhibitory activities against these CDKs could confound the assessment of Q901’s effects on transcription via CDK7 inhibition, as noted in prior studies with other CDK7 inhibitors such as THZ1 (*12–14*). The IC_50_ values of Q901 were found to be greater than 10 μM for CDK8, 8.58 μM for CDK9, 2.48 μM for CDK12, and 7.69 μM for CDK13. The fold differences in Q901-mediated inhibition of these CDKs compared to CDK7 were >1,000 for CDK8, 715 for CDK9, 207 for CDK12, and 641 for CDK13 (Fig. 1D, and table S1). These analyses collectively highlight the superior selectivity and efficacy of Q901 as a CDK7 inhibitor compared to previously characterized inhibitors (*12–14*).

### Q901 alters RNAPII transcription profiles

Various cancer tissues, including estrogen receptor-positive (ER+) breast cancer often exhibit a higher expression levels of CDK7 compared to their normal counterparts, which are also associated with poorer patient outcomes (*9*). MCF7 cells, a representative ER-positive breast cancer model, also exhibit high expression levels of CDK7 (*11*). Inhibition of CDK7 has been shown to suppress cell cycle progression and induce apoptosis in MCF7 cells, suggesting that targeting CDK7 could be a viable therapeutic strategy for breast cancer (*26*). For this reason, we used ER-positive MCF-7 cells as a model system to evaluate the efficacy of Q901. Q901 exhibited effective inhibition of MCF-7 cell growth with IC_50_ of 95 nM at 72 h, 16 nM at 96 h, and 5 nM at 120 h of treatment (Fig. 2A).

**Fig. 2.**
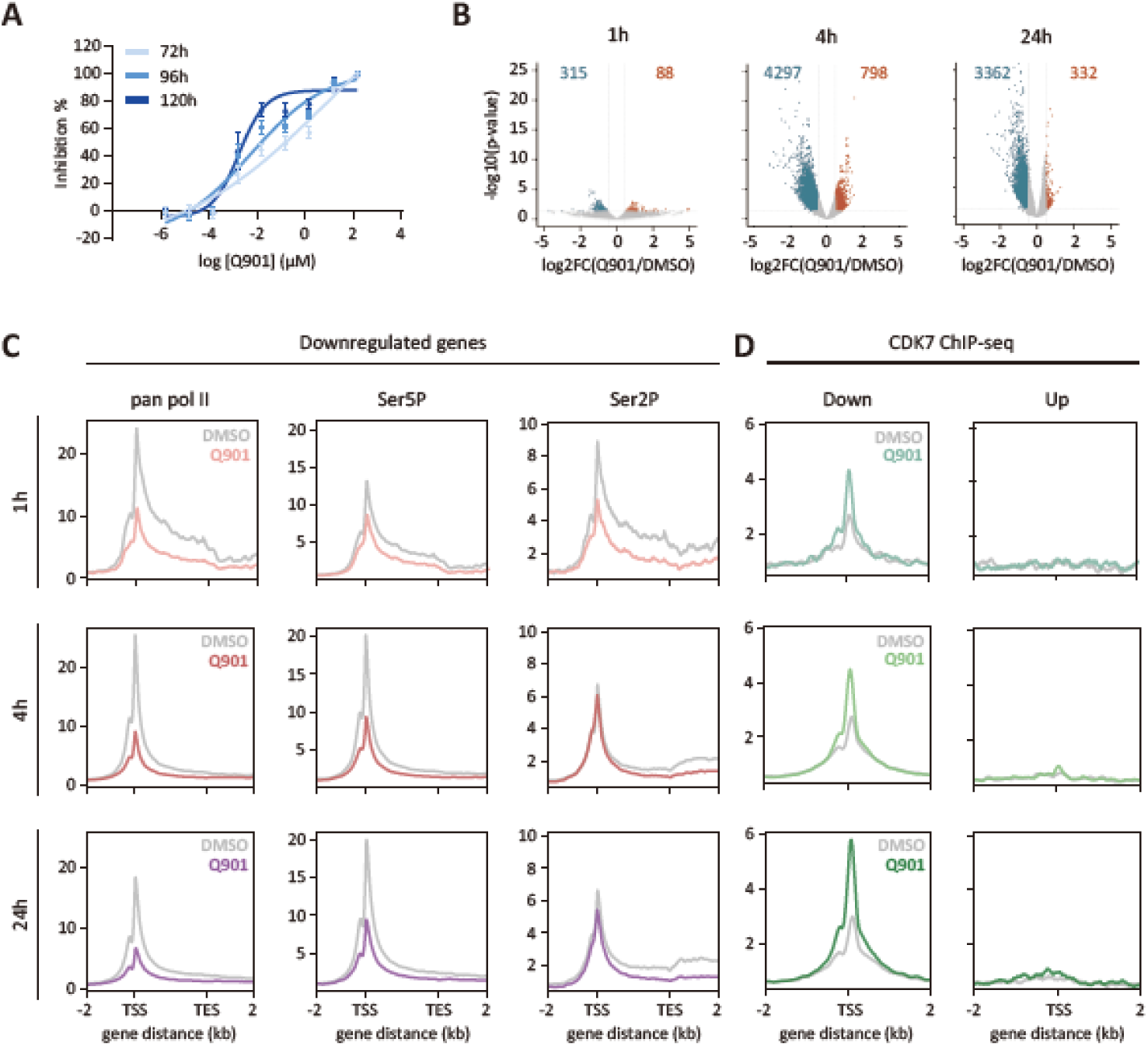
Q901 alters RNAPII phosphorylation and transcriptional activity. (**A**) MCF-7 cells were treated with Q901 at the indicated concentrations for 72, 96, or 120 h. Cell viability was measured using the ATP Lite™ system. Inhibition (%) was plotted against the log-transformed Q901 concentration (µM) (n = 3 to 4). Data represent mean ± SD. (**B** and **C**) RNAPII ChIP-seq was performed following treatment with 100 nM Q901 for the indicated duration. (B) Volcano plot of pan RNAPII ChIP-seq signals after Q901 treatment (n = 2; blue dot indicates p-value ≤ 0.05 and log2FC ≤ -0.58; red dot indicates p-value ≤ 0.05 and log2FC ≥ 0.58). (C) Average ChIP-seq signal plots of various RNAPII forms for genes downregulated by Q901. (**D**) Average CDK7 ChIP-seq signal plots of downregulated and upregulated genes at the TSS. CDK7 ChIP-seq was performed with 100 nM Q901 for the indicated durations.

To assess the phosphorylation levels of the CTD in RNAPII bound to chromatin, ChIP-seq experiments were conducted using specific antibodies targeting individual phosphorylated serine residues (Fig. 2, B and C). Volcano plot analysis revealed reduced RNAPII binding in thousands of genes following Q901 treatment, with the magnitude of reduction increasing over time (Fig. 2, B and C, and fig. S2A). Hundreds of genes including the gene encoding the largest subunit of RNAPII (POLR2A) a biomarker for pharmacodynamic (PD) responses to CDK7 inhibitors (*15, 27*) exhibited a slight increase in RNAPII binding (Fig. S2, B and C). However, the average level of pan RNAPII in these upregulated genes was significantly lower than that in downregulated genes. Additionally, a slight increase in RNAPII levels was observed only at the 1 and 4 h time points after Q901 treatment, but not at 24 h. This suggests that the observed increase in RNAPII binding may be a temporary effect resulting from the global downregulation of RNAPII binding to chromatin as part of a compensatory mechanism. In contrast, downregulated genes exhibited significantly higher levels of basal transcription. Analysis of their mRNA half-lives showed a trend toward a negative correlation between the degree of transcriptional downregulation and mRNA stability; specifically, log2 fold change (Q901/DMSO) values were relatively higher in mRNAs with shorter half-lives (Fig. S2D). This finding aligns with a previous study indicating that CDK7 inhibition in A549 cells primarily downregulated short-lived genes with high mRNA turnover rates (*2*).

The occupancy levels of Ser2 and Ser5 phosphorylated forms of RNAPII were also significantly decreased upon Q901 treatment (Fig. 2C). The decrease in Ser5 phosphorylation levels most likely resulted from the direct inhibition of CDK7 by Q901. Unlike the Ser5 phosphorylation profiles, which showed strong enrichment near the transcription start sites (TSSs), the decrease in Ser2 phosphorylation was more prominent in the gene coding region, especially toward the transcription end sites (TESs) at later time points of Q901 treatment. This result was expected, as CDK9, a primary kinase that phosphorylates Ser2 during the elongation phase, needs to be activated by CDK7 through phosphorylation (*6*).

Q901 also altered the binding profile of CDK7 at the promoter regions (Fig. 2D). TFIIH associates with the promoter to form a preinitiation complex with RNAPII, facilitating the transition from initiation to elongation (promoter clearance). Following this, it is released from early elongation complexes and recycled for subsequent rounds of transcription (*7*). However, Q901 treatment significantly increased CDK7 occupancy over time, indicating that covalently inhibited CDK7 remained stably bound to the promoter without being released (Fig. 2D left). This phenomenon was not observed in upregulated genes. In fact, no significant CDK7 signals were detected at the promoters of upregulated genes, regardless of Q901 treatment (Fig. 2D right). It remains unclear whether the upregulated genes can circumvent CDK7 action during the transcription cycle or if the observed results are merely due to detection sensitivity issues stemming from the significantly lower levels of basal transcription in this gene group.

Taken together, our findings support a model in which the inhibition of CDK7 activity alters the dynamics of transcription initiation complexes, as previously described (*2, 6, 28, 29*). Given CDK7’s established role in RNAPII pausing (*6, 8, 28, 29*), the reduced occupancy of RNAPII at promoter-proximal regions upon Q901 treatment likely results from the rapid clearance of RNAPII complexes from their pausing sites. Furthermore, a recent study has shown that CDK7 activity is crucial for the efficient release of transcription initiation factors and the Mediator complex (*28*). In this context, the observed Q901-mediated trapping of CDK7 could lead to the prolongation of the RNAPII preinitiation complex at the promoters, preventing transcription factor cycling.

### Q901 impairs enhancer function

Super-enhancers (SEs) are clusters of enhancers that control genes critical for cellular identity and function (*30*). SEs are densely enriched with transcription factors, RNAPII machinery, and chromatin regulators, driving high transcription levels of enhancer RNAs (eRNAs) as well as mRNAs from their target genes (*31*). Dysregulation of SE activity is common in cancer, functioning as a major oncogenic driver. In such cases, cancer cells become transcriptionally addicted, requiring higher levels of transcription activity than normal cells to maintain their growth. This transcriptional addiction provides a therapeutic opportunity to target the general transcription machinery to reduce transcriptional output (*9*).

For this reason, we investigated the effect of Q901 on SE-driven transcriptional activities. In MCF-7 cells, 483 SEs and 8,134 typical enhancers (TEs) were identified using publicly available H3K27ac ChIP-seq data and the ROSE program (Fig. S3A) (*32*). The identified enhancers were classified into four groups: CDK7-bound SEs, CDK7-nonbound SEs, CDK7-bound TEs, and CDK7-nonbound TEs. Both SEs and TEs were strongly enriched with CDK7 and RNAPII, with SEs exhibiting stronger enrichment than TEs (Fig. 3A). A similar pattern was observed for enhancer RNA profiles, reflecting a high level of transcription activity at enhancers (Fig. S3C).

**Fig. 3.**
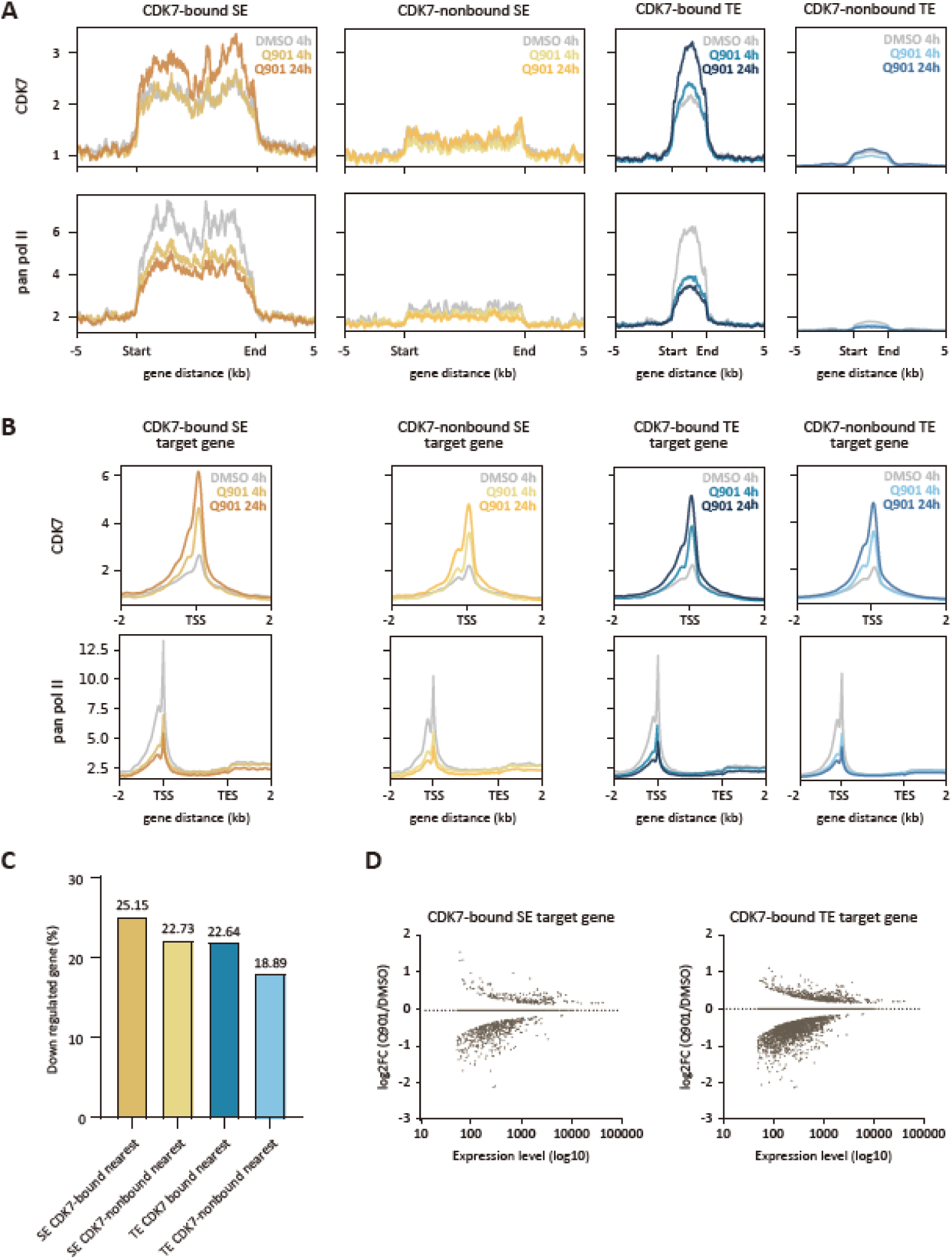
Q901 reduces enhancer-associated transcriptional activity. (**A**) Average ChIP-seq signal plots of CDK7 and pan RNAPII ChIP-seq across four enhancer groups. (**B**) Average ChIP-seq signal plots for CDK7 and pan RNAPII for protein-coding genes regulated by four enhancer groups. (**C**) Bar graph shows the proportion of downregulated genes in each enhancer groups (Pan RNAPII ChIP-seq; n = 2; p-value ≤ 0.05 and log2FC ≤ -0.58). (**D**) Scatter plot shows the expression levels and log2FC of target genes regulated by CDK7-bound SE and CDK7-bound TE (pan RNAPII ChIP-seq; n = 2; p-value ≤ 0.05).

Q901-mediated inhibition of CDK7 activity significantly reduced RNAPII binding at SEs and TEs, suggesting that Q901 impaired enhancer activities (Fig. 3A, and fig. S3C). To examine the impact of enhancer inhibition on target gene transcription, RNAPII and CDK7 binding levels were assessed in genes located within ± 50 kb from the enhancer regions (Fig. 3B). Baseline RNAPII binding was highest in the target genes of CDK7-bound SEs (Fig. S3B). The proportion of downregulated genes was also highest in that group (∼25% of potential target genes) (Fig. 3C). However, the degrees of transcriptional alteration caused by Q901 were comparable across the four enhancer target gene groups, with no strong bias toward SE-regulated genes (Fig. 3, C and D).

Notably, Gene Ontology (GO) analysis revealed that genes involved in nucleosome organization were primarily affected by CDK7 inhibition in SEs, whereas the downregulation of genes related to translation resulted from alterations in TE activities (Fig. S3D). These findings suggest that the anticancer effect of Q901 is, in part, mediated by suppressing the transcription of genes necessary for cell growth and proliferation through modulation of enhancer activities.

### CDK7 inhibition impairs key oncogenic pathways

To further elucidate the molecular consequences of CDK7 inhibition, we performed Gene Set Enrichment Analysis (GSEA) utilizing hallmark gene sets (Fig. 4A). Downregulated genes were significantly enriched in pathways canonically associated with oncogenesis, including cell cycle control, DNA repair mechanisms, and targets of the oncogenic transcription factors (TFs) MYC and E2F. CDK7 occupancy at these pathway genes was markedly elevated over time with Q901 treatment, whereas RNAPII levels were reciprocally diminished, substantiating the efficacy of Q901 as a CDK7 inhibitor (Fig. S4, A to E). Corroborating the pathway analysis, MYC and E2F ChIP-seq revealed attenuated binding of these TFs at target gene promoters after treatment of Q901 (Fig. 4, B and C).

**Fig. 4.**
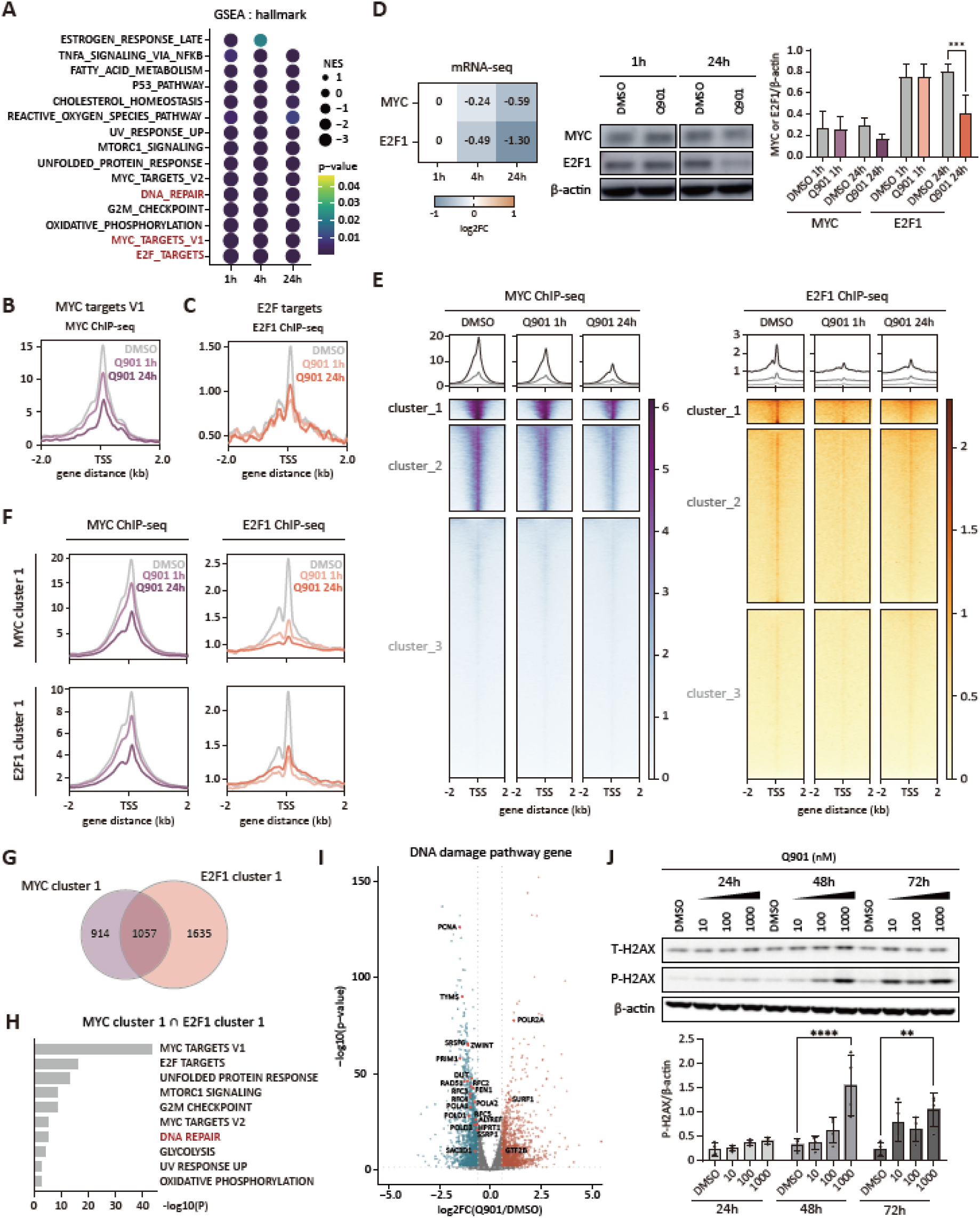
Q901 disrupts MYC-E2F-DNA damage repair transcriptional axis. (**A**) GSEA of pan RNAPII ChIP-seq data following Q901 treatment (n = 2; fgsea p-value ≤ 0.05). (**B**) Average MYC ChIP-seq signal plot of MYC targets V1 gene set. (**C**) Average E2F1 ChIP-seq signal plot of E2F targets V1 gene set. (**D**) Heatmap showing log2FC values of MYC and E2F1 expression from mRNA-seq data and western blot showing MYC and E2F1 protein levels with or without Q901 treatment (right; n = 3; two-way ANOVA followed by Šidák’s multiple comparison, data represent mean ± SD; ***: p < 0.001). (**E**) Average plots and heatmaps showing MYC ChIP-seq signals for MYC-clustered genes (left), and E2F1 ChIP-seq signals for E2F1-clustered genes (right). (**F**) Average plots of MYC and E2F1 ChIP-seq signals for MYC cluster 1 and E2F1 cluster 1 genes. (**G**) Venn diagram showing the overlap between MYC cluster 1 and E2F1 cluster 1 protein-coding genes. (**H**) GO analysis of protein-coding genes commonly present in MYC cluster 1 and E2F1 cluster 1. (**I**) Volcano plot of mRNA-seq signals after Q901 24h treatment (n = 3; blue dot indicates p-value ≤ 0.05 and log2FC ≤ -0.58; red dot indicates p-value ≤ 0.05 and log2FC ≥ 0.58). DNA damage pathway genes indicated by a red dot and labeled. (**J**) Western blot data of total H2AX and its phosphorylated form of H2AX (γH2AX) after Q901 treatment for indicated times (n = 4; ordinary one-way ANOVA followed by Šidák’s multiple comparison, data represent mean ± SD; **: p < 0.01, ***: p < 0.001).

To investigate the mechanism by which CDK7 inhibition modulates the binding properties of MYC and E2F, we quantified changes in their expression levels at 1 and 24 h post-Q901 treatment (Fig. 4D). Only E2F protein level showed a significant reduction at the 24 h time point (Fig. 4D middle and right). Therefore, the diminished binding of both TFs, observed at a much earlier time point (1 h) following Q901 administration, may not be solely ascribed to altered protein levels. Consistently, we observed no alterations in the phosphorylation status of MYC at residues S62 and T58, which have been previously implicated in MYC protein stability (Fig. S5A) (*33*). The retinoblastoma protein (Rb) is known to exert cell cycle suppression through direct interaction with E2F, thereby sequestering it (*34*). Phosphorylation of Rb at these specific residues triggers the dissociation of E2F, facilitating the transcriptional activation of genes essential for cell cycle progression. However, Q901 treatment did not induce significant changes in Rb phosphorylation at residues S780 and S795, suggesting that changes in Rb-mediated E2F sequestration are unlikely to cause the observed reduction in E2F binding to target gene promoters (Fig. S5A). We also found no significant changes in protein levels of MYC-dimerization partner, MAX (Fig. S5B). Therefore, inhibition of key oncogenic pathways by Q901 appears to be mediated by both altering the TF binding properties and the expression levels during the course of CDK7 inhibition.

### CDK7 inhibition impairs DNA repair pathway by controlling MYC and E2F1 activity

MYC and E2F are key TFs involved in cell cycle progression, cell growth and proliferation, which are crucial for cancer progression and survival. To gain additional molecular insight into the effect of Q901-mediated CDK7 inhibition in MYC and E2F activity, the entire genes were ranked based on MYC or E2F1 occupancy levels at the TSS regions, and the highest occupancy cluster (cluster 1) for each TF was compared for Q901 effects (Fig. 4E). Both TFs largely co-occupied the genes at each cluster 1 and exhibited reduced occupancy upon Q901 treatment (Fig. 4, F and G), suggesting an extensive co-operation of MYC and E2F1 in MCF-7 cells. Both TFs were also enriched at enhancers (Fig. S6). Most SEs were co-bounded by both TFs, in contrast to TEs where only approximately half displayed co-binding (Fig. S6A). Q901 significantly reduced TF occupancy levels at enhancers as well as their target genes (Fig. S6, B and C). Our analysis collectively suggests that Q901 could suppress the MYC and E2F-driven oncogenic transcription program by reducing their binding activities at both promoters and enhancers.

Interestingly, GO analysis with the co-occupied cluster 1 genes revealed several enriched pathways that were also significantly altered upon CDK7 inhibition in the GSEA using the entire genes, indicating that the Q901 effect was mainly on genes regulated by MYC and E2F (Fig. 4, A and H). We focused on DNA damage repair (DDR) gene set since the accumulation of the DNA damage has been well established as a key mechanism in anticancer activity (*35*). We observed that most DDR genes were downregulated following Q901 treatment (Fig. 4I, and fig. S4F). Since downregulation of DDR genes by Q901 could lead to accumulation of DNA damage, we examined DNA damage marker, histone γH2AX phosphorylation level and found that there was an increase in γH2AX phosphorylation level over time in a dose-dependent manner (Fig. 4J). Therefore, Q901 can lead to DDR deficiency in MCF-7 cells by altering transcription activity of MYC and E2F at both promoters and enhancers, and subsequent downregulation of genes involved in the DNA damage repair pathway.

### Q901 disrupts the TOP1-MYC complex at the TSS and extends the stability of TOP1-DPCs

A prominent function of MYC in cancer is transcriptional amplification. Instead of controlling a specific set of genes, MYC primarily contributes to the amplification of transcription outputs of active genes (*36*). A high level of transcription activity can cause significant DNA topological stress through the formation of negative and positive supercoiling at downstream and upstream of elongating RNAPII, respectively, which would eventually halt transcription. MYC has been shown to overcome torsional stress during transcription by recruiting and activating TOP1 and topoisomerase II (TOP2) at promoters, gene bodies, and enhancers (*37*). As MYC binding to the TSSs and the enhancers was impaired by Q901-mediated CDK7 inhibition, we predicted that the TOP1 recruitment might be impaired at MYC-bound target genes by Q901. Thus, we performed TOP1 ChIP-seq to examine the impact of Q901 on TOP1 binding. The binding of TOP1 at MYC target genes correlated directly with MYC binding levels, and treatment with Q901 resulted in decreased TOP1 binding at these genomic sites without altering overall TOP1 expression (Fig. 5A, and fig. S7, A and B).

**Fig. 5.**
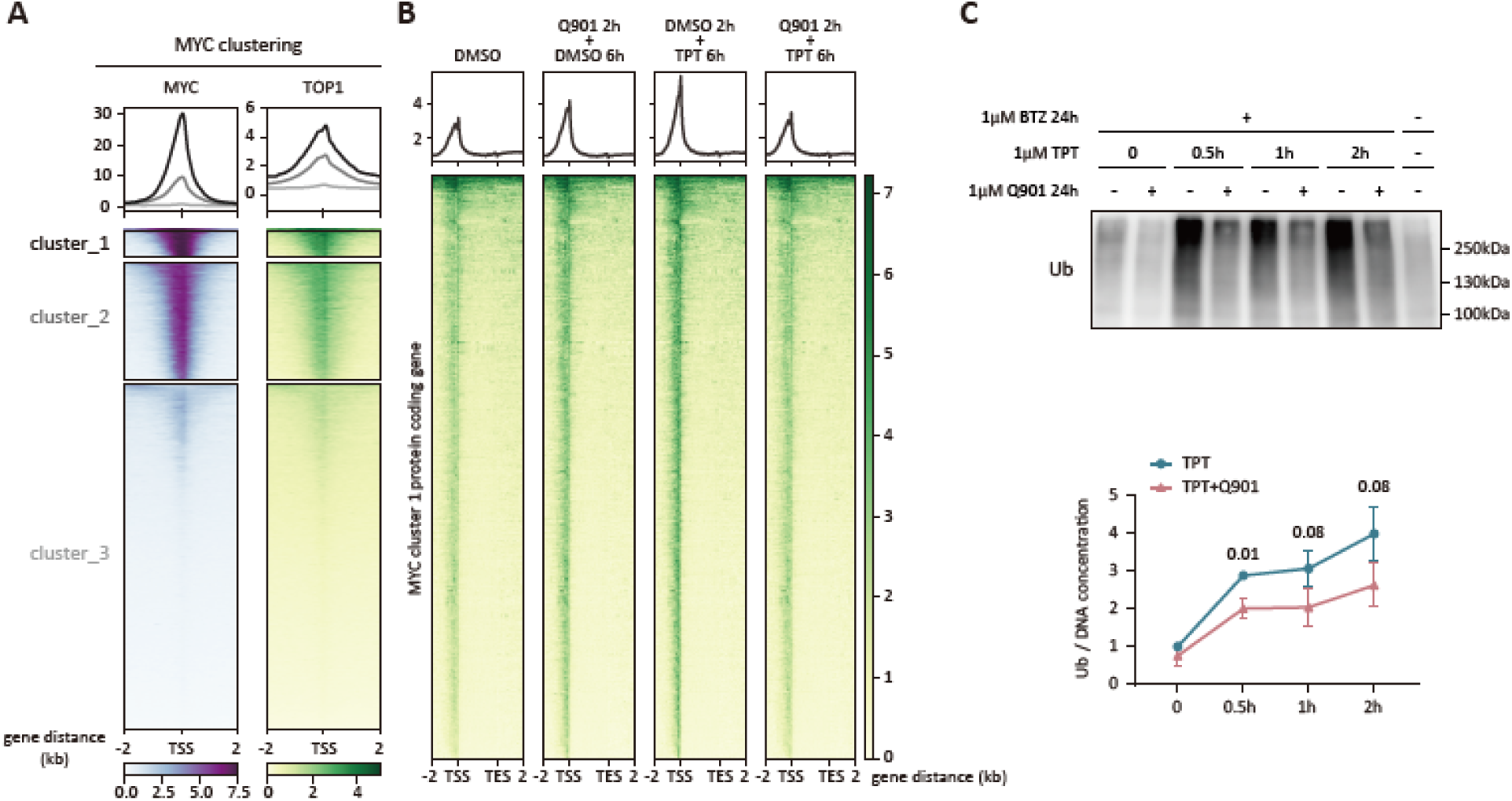
Q901 reduces the TSS-associated TOP1-MYC complex and prolongs TOP1-DPC stability. (**A**) Average plots and heatmaps of MYC and TOP1 ChIP-seq signals at three gene clusters ranked by MYC binding levels (DMSO condition). (**B**) Average plots and heatmaps of TOP1 ChIP-seq signals for MYC cluster 1 protein-coding genes after Q901 or TPT treatment. (**C**) Detection of ubiquitinated TOP1-DPCs by DUST assay (top), with quantification of Ub-TOP1-DPC levels (bottom) (n = 3; multiple unpaired t-test followed by two-stage step-up method of Benjamini, Krieger and Yekutieli, data represent mean ± SD).

We next examined the effect of TOP1 inhibitor, TPT which can trap the TOP1ccs, generating TOP1-DPCs (*18*). TOP1 ChIP-seq in the presence of TPT showed increased binding of TOP1 at MYC target genes, likely representing the TOP1-DPCs (Fig. 5B). TOP1-DPCs represent a unique type of DNA lesion that can arrest RNAPII elongation complexes. When RNAPII becomes arrested at a TOP1-DPC, it triggers ubiquitination and 26S proteasome-mediated degradation of both RNAPII and TOP1-DPC, which activates transcription-coupled DNA repair (TCR) (*17*). If these TOP1-DPCs remain unresolved, they can ultimately lead to cell death (*17, 18*). A recent study in small-cell lung cancer cells has shown that THZ1, another CDK7 inhibitor, enhanced the anticancer activity of TPT by promoting ubiquitin-mediated degradation of RNAPII, resulting in unresolved TOP1-DPCs (*25*). Based on our finding that CDK7 inhibition by Q901 decreased RNAPII at the promoter-proximal regions (Fig. 2C), we hypothesized that Q901 could also synergize the activity of TPT by decreasing the population of elongating RNAPII encountering TOP1-DPCs, which would otherwise trigger proteasomal degradation of RNAPII and TOP1-DPC. Consistently, TOP1 ChIP-seq result showed that Q901 counterbalanced the TPT effect in TOP1-DPC stability at MYC target genes when MCF-7 cells were treated with both (Fig. 5B).

To further demonstrate this effect, DUST (Detection of Ubiquitylated and SUMOylated TOP1-DPC) assay was performed to examine the effect of Q901 in TPT-mediated ubiquitination of TOP1-DPCs (Fig. 5C, and fig. S7C). MCF-7 cells were treated with either TPT alone or TPT+Q901 at various times in the presence of protease inhibitor, Bortezomib (BTZ), and the amounts of ubiquitinated TOP1-DPC were measured by western blot using ubiquitin (Ub)-specific antibody. TPT alone-treated cells exhibited higher levels of TOP1-DPC ubiquitination at all time points tested, than TPT+Q901-treated cells (Fig. 5C, and fig. S7C).

Taken together with the results shown in Fig. 2C, our findings provide a mechanistic basis for combining Q901 with TPT as a therapeutic approach in cancer treatment. Q901-mediated CDK7 inhibition blocks the transition of RNAPII into stable elongation, thereby decreasing RNAPII levels at MYC target genes. This reduction in elongating RNAPII has an important consequence: fewer RNAPII complexes encounter and become arrested at TOP1-DPC sites generated by TPT. Since arrested RNAPII complexes typically activate the proteasome pathway to remove TOP1-DPCs, the Q901-mediated decrease in elongating RNAPII results in fewer clearance events, allowing TOP1-DPCs to persist longer in cancer cells.

### Combination of Q901 and topoisomerase I inhibitors induces synergistic anti-tumor effect

Our study demonstrates that Q901-mediated CDK7 inhibition synergizes with TOP1 inhibitors through three distinct mechanisms: (1) reducing MYC and E2F recruitment to oncogenic transcription programs, (2) impairing DNA damage repair pathways by downregulating DDR genes, and (3) prolonging TOP1-DPCs by blocking their proteasomal degradation through RNAPII displacement (Fig. 6A).

**Fig. 6.**
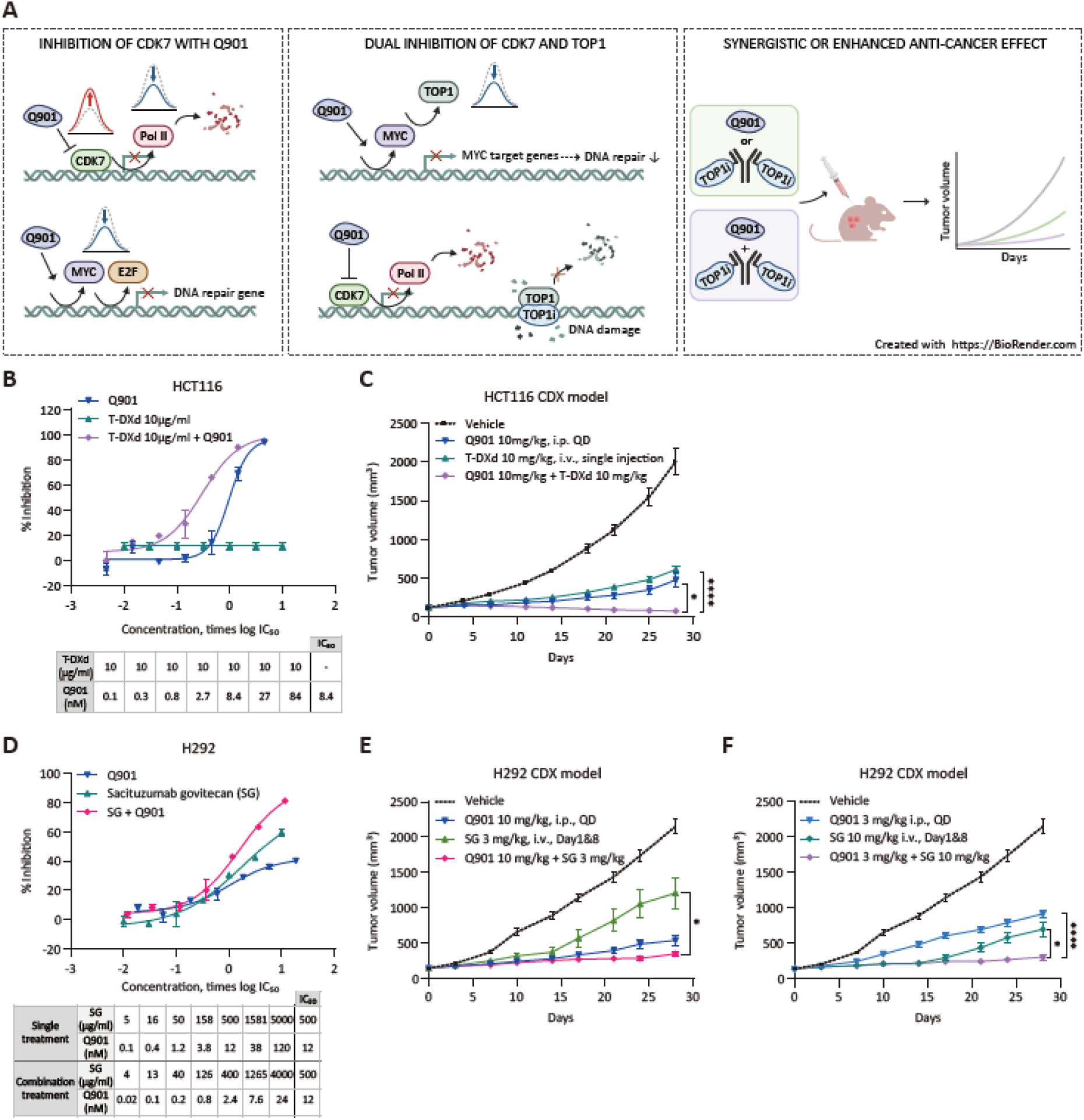
Q901 synergies with TOP1i-ADCs. (**A**) A model illustrating how Q901 enhances the activity of TOP1i and TOP1i-ADCs. (Left) Q901 promotes CDK7 accumulation at the TSS while reducing RNAPII binding. This also decreases MYC and E2F1 binding at the TSS, leading to downregulation of genes involved in the DNA damage response pathway. (Middle) The dual inhibition of CDK7 (by Q901) and TOP1 (by TOP1i) blocks the repair of TOP1i-induced DNA damage, ultimately leading to cell death. (Right) The combination of Q901 and a TOP1i-ADC shows potent enhanced anticancer activity, effectively inducing cancer cell death in vitro and significantly reducing tumor growth in vivo. (**B** and **C**) HCT116, HER2 ultra low/negative human colon cancer cell line, was treated with Q901, T-DXd (10 μg/ml), or their combination at the indicated concentrations for 72 h (B). Dose-response curves were plotted as a function of log-transformed concentration relative to IC□□ values. Cell viability was measured using the ATP Lite™ system (n = 2, data represent mean ± SD). (C) For in vivo efficacy study, HCT116 cells mixed with Matrigel (1:1) were subcutaneously implanted into BALB/c nude mice. When tumors reached an average size of 117 mm3, mice were randomized into groups (n = 8 per group) and treated with Q901 (10 mg/kg, intraperitoneally once daily), T-DXd alone (10 mg/kg, intravenous single injection on day 0) or the combination. (**D** and **F**) H292, TROP2 positive human lung cancer cell line, was treated with Q901 in combination with SG at the indicated concentrations for 72 h (D). Dose-response curves were generated using log-transformed concentrations normalized to IC□□ values. Cell viability was assessed using the ATP Lite™ system (n = 2, data represent mean ± SD). (**E** and **F**) For in vivo efficacy study, H292 cells were mixed with Matrigel (1:1) and implanted subcutaneously into BALB/c nude mice. When tumors reached an average size of 140 mm3, mice were randomized into groups (n = 8 per group) and treated with Q901 (10 or 3 mg/kg, intraperitoneally once daily), SG alone (3 or 10 mg/kg, intravenous single injection on day 1 and 8) or the combination of both. The graph shows the mean tumor volume ± SEM. Statistical significance was calculated using GraphPad Prism software (*: p < 0.01, ****: p < 0.0001 by two-way ANOVA followed by Tukey’s multiple comparison test).

To validate these synergistic mechanisms across multiple cancer models, we first examined rapidly proliferating neuroendocrine small cell lung cancer (SCLC) cell lines, H82 and DMS53. SCLC was chosen as a clinically relevant model for testing synergistic mechanisms, particularly given that TOP1 inhibitors such as TPT, are already utilized in clinical settings for SCLC treatment. As expected, the combined administration of Q901 and TPT with Q901 and TPT for 72 h resulted in enhanced antiproliferative activity compared to either drug alone (Fig. S8, A and B). We next examined whether Q901 could enhance the efficacy of T-DXd, a HER2-targeting ADC carrying a TOP1 inhibitor payload. Because cellular uptake of the TOP1 inhibitor component of T-DXd depends on HER2 abundance (20, 22, 38), we focused on a system with low HER2 expression to assess the combination’s efficacy under unfavorable delivery conditions. We selected the HCT116 colon cancer model, which exhibits ultra-low HER2 expression and shows no response to T-DXd in cellular cytotoxicity assays, likely due to insufficient intracellular delivery of DXd (Fig. 6B). While these cells were insensitive to T-DXd monotherapy (10 μg/ml), Q901 alone inhibited cell growth with an IC_50_ of 8.4 nM. The combination of T-DXd and Q901 was assessed based on the combination synergy finding workflow suggested by Uitdehagg et al. (*38*). The test essentially examines various combination ratios of two drug to see if there is any synergy by combining two drugs. Importantly, the combination of highest concentration (10 μg/ml) of T-DXd (inactive) with various Q901 concentration ratios compared to Q901 IC_50_ in HCT116 cell line significantly enhanced growth inhibition across most dose ratios tested, demonstrating that Q901 can synergize with a low dose of DXd that cannot exert any cytotoxic effect by itself.

Next, the *in vivo* antitumor activity of this combination was evaluated in HCT116 xenograft models. Mice bearing HCT116 tumors were treated with T-DXd (10 mg/kg), Q901 (10 mg/kg), or their combination (Fig. 6C). By day 28, T-DXd and Q901 monotherapies achieved similar tumor growth inhibition (TGI) of 74% and 81%, respectively. Notably, despite the lack of response to T-DXd in cellular assays, substantial tumor growth inhibition was observed in vivo. This discrepancy is likely attributable to systemic release of DXd payload from circulating T-DXd, leading to free payload-mediated antitumor effects (*39*). To test this possibility, T-Dxd was incubated in the mouse serum for 21 days and measured drug-to-antibody ratio (DAR). The results showed a marked decrease in DAR values by day 21, indicating that extensive Dxd payload release into the mouse serum is a likely cause of the discrepancy between the cellular assay and the mouse xenograft test (Fig. S9). Nevertheless, the combination treatment enhanced the efficacy of tumor suppression with 102% TGI. While tumors in the monotherapy groups resumed growth approximately 11 days post-treatment and gradually increased in size, the Q901/T-DXd combination exhibited tumor suppression throughout the study duration.

To further extend our findings, we investigated another TOP1i-ADC, sacituzumab govitecan (SG), which targets trophoblast cell surface antigen 2 (TROP2)—a transmembrane glycoprotein frequently overexpressed in many epithelial malignancies and associated with aggressive tumor behavior. SG binds to TROP2 on the tumor cell surface, undergoes receptor-mediated internalization, and is cleaved intracellularly to release SN-38, a potent TOP1 inhibitor that induces DNA damage in tumor cells (41, 42). Because clinical outcomes of SG treatment have been reported to be independent of TROP2 expression levels, we selected a TROP2-positive cell line to evaluate the combination effect of Q901 under conditions where SG is expected to be active. In TROP2-positive H292 lung cancer cells, both SG (IC_50_ 500 ng/ml) and Q901 (IC_50_ 12 nM) as monotherapies inhibited cell proliferation, but their combination significantly increased the antiproliferative effect (Fig. 6D). In H292 xenograft models, two dosing regimens were evaluated. In the first, SG monotherapy at 3 mg/kg achieved 47% TGI and Q901 monotherapy at 10 mg/kg achieved 80% TGI by day 28. The combination treatment enhanced the efficacy to 90% TGI (Fig. 6E). In the second regimen, SG monotherapy at 10 mg/kg achieved 72% TGI and Q901 monotherapy at 3 mg/kg achieved 61% TGI by day 28. The combination treatment enhanced efficacy to 92% TGI (Fig. 6F). Notably, tumors treated with SG alone showed regrowth after 14 days, while both Q901 monotherapy and the combination treatment maintained sustained tumor growth inhibition throughout the study period, with the combination showing slightly higher efficacy. Collectively, these results highlight Q901 as a promising combination partner for topoisomerase I inhibitors and TOP1i-ADCs across multiple cancer models. Q901 not only enhances efficacy in tumors already responsive to TOP1i-ADCs, but also provides benefit in setting with limited ADC activity, such as those with low antigen expression or reduced TOP1 inhibitor exposure due to suboptimal dosing.

## Discussion

Our study presents Q901 as a highly selective CDK7 inhibitor with potential to address challenges in cancer therapy, particularly in combination with TOP1 inhibitors. By targeting CDK7, Q901 traps CDK7 at promoter-proximal regions, preventing RNAPII from progressing into elongation. In MCF-7 cells, CDK7 inhibition primarily disrupts oncogenic transcription programs driven by MYC and E2F and impairs the DNA damage response pathway. These effects enhance tumor sensitivity to TOP1 inhibitors by prolonging DNA damages caused by TOP1-DPCs. These findings underscore the therapeutic promise of Q901, especially in cancers with high transcriptional activity and limited response to current TOP1 inhibitor-based therapies.

Preclinical studies of CDK7 inhibitors have demonstrated anticancer activities through cell cycle arrest and suppression of transcription (*12–14*). While cell cycle arrest was commonly observed with all tested inhibitors, their effects in transcription and mechanisms of action varied depending on the type of inhibitor used. THZ1 and its derivatives (SY-1365 and SY-5609) induce G2/M cell cycle arrest and apoptosis, exhibiting strong anticancer effects against various cancer types (*13*). Global transcription repression was also observed, accompanied by a significant loss of RNAPII CTD phosphorylation (*12, 13, 15*). The extreme sensitivity of T-cell leukemia (T-ALL) cells to THZ1 was attributed to transcription addiction, especially those genes controlled by super-enhancers and often driven by oncogenic transcription factors like MYC (*13*). However, the observed anticancer effects through transcriptional inhibition were later found to be contributed by off-target inhibition of CDK12/13, rather than through CDK7 inhibition alone (*12, 13, 15*).

In contrast, YKL-5-124, a highly selective CDK7 inhibitor with no measurable affinity for CDK12/13, showed transcriptional repression in rather selected gene expression programs, including E2F and MYC pathways, without broadly affecting super-enhancer-mediated or global transcription. Treatment with YKL-5-124 did not result in significant reductions in bulk RNAPII phosphorylation and primarily induced G1 phase cell cycle arrest without noticeable evidence of apoptosis (*14*). These findings collectively suggested a non-essential role for CDK7 in regulating RNAPII CTD phosphorylation and global basal gene expression. However, it causes DNA replication stress and genomic instability, which in turn activate antitumor immune responses by promoting lymphocyte activation (*40*).

Q901 exhibits high selectivity for CDK7 over other CDKs (Fig. 1). Its anticancer effects are being confirmed in the preclinical as well as in the clinical studies. Exhibiting no off-target effects in CDK12/13 (Fig. 1), the selectivity of Q901 appears to be similar to that of YKL-5-124, yet our study delineates more defined transcriptional mechanism elicited by specific CDK7 inhibition, some of which had not been previously appreciated. Q901 significantly decreased the levels of chromatin-bound RNAPII and CTD phosphorylation at thousands of genes (Fig. 2, B and C). Notably, catalytically inhibited CDK7 became trapped at the promoter-proximal regions (Fig. 2D). These findings are in line with a recent study demonstrating the direct role of CDK7 in RNAPII promoter escape by facilitating initiation factor release (*28*). Rapid CDK7 inhibition caused RNAPII retention at promoters, leading to decreased RNAPII initiation and global downregulation of transcription. We envision that trapping of catalytically inhibited CDK7 appears to be the key mechanism for the retention of RNAPII initiation complex at the promoter-proximal regions. Q901 also altered the enhancer activity by reducing RNAPII binding and eRNA transcription at both SEs and TEs, leading to impaired transcription from their predicted target genes. In contrast with the preferential inhibitory activity of THZ1 on oncogenic programs driven by SEs, there was no bias in the effect of Q901-mediated CDK7 inhibition toward altering SE activities compared to TEs in MCF-7 cells (Fig. 3). Nonetheless, enhancer target genes affected by Q901 were enriched in pathways related to nucleosome organization and translation, suggesting that the anticancer effect of Q901 is, in part, mediated by suppressing enhancer activities (Fig. S3D).

Our systematic analysis identified that MYC and E2F occupied the promoters of nearly all expressed genes in MCF-7 cells, with their occupancy levels being highly correlated with transcription levels (Fig. 4E). Q901 globally decreased their binding levels (Fig. 4, E and F). MYC can promote transcription through multiple mechanisms such as facilitation of the RNAPII pause-release and the recruitment and activation of TOP1 to alleviate DNA topological stress during active transcription (*37*). Therefore, attenuated MYC binding would further contribute to the anticancer activity of Q901.

ADCs represent a rapidly expanding class of anti-cancer therapeutics, with 14 FDA-approved agents on the market. TOP1 inhibitors are increasingly used as payloads to enhance both the therapeutic index and targeted delivery to antigen-expressing tumors, as they generally exhibit manageable toxicity compared to other anticancer agents while remaining highly effective in cancers that require high levels of transcription and replication (*19*). Besides already approved T-DXd (Enhertu) and SG, more than 100 TOP1i-ADC candidates are currently in clinical trials (*41*). Despite their promising potential, the development of new TOP1i-ADCs faces significant challenges, primarily due to two factors: 1) the need for an increased therapeutic window, and 2) the necessity of overcoming cross-resistance issues when treating patients with ADCs in sequences. Toxicities of TOP1i-ADCs often resembles those of the standalone TOP1 inhibitor chemotherapies. To improve the toxicity profile of ADCs, multiple strategies are being employed, especially through advancements in ADC engineering such as site-specific conjugation, the use of stable linkers, and hydrophobicity masking technologies (*42*). However, these engineering approaches have yet to demonstrate significant clinical benefits (*41*). With numerous ADCs are under clinical development, there is a concern of cross-resistance when a cancer patient is treated with sequential use of ADCs (*41*). Although limited, some clinical data suggest that activation of DNA damage repair may be a common mechanism driving resistance to sequential TOP1i-ADC treatments (*43, 44*). Our study identified Q901 as a promising combination partner for TOP1 inhibitors and TOP1i-ADCs across multiple preclinical cancer models (Fig. 6A). Mechanistically, Q901-mediated CDK7 inhibition suppressed DDR pathways by transcriptionally regulating MYC and E2F activity (Fig. 4). Additionally, it prevented the proteasomal degradation of TOP1-DPCs by disrupting the transition of RNAPII complex into elongation (Fig. 5).

Our study strongly supports the combinatorial use of Q901 and TOP1i-ADCs as a novel strategy to enhance ADC efficacy and broaden their therapeutic potential in cancer treatment. This approach may be particularly beneficial in settings where ADC activity is limited, such as in tumors with low antigen expression that restricts TOP1 inhibitor exposure, when dose reductions are required to manage toxicity, or when multiple TOP1i-ADCs are administered sequentially.

## Materials and Methods

### Cell lines and reagents

MCF-7, H82, HCT116, and H292 cell lines were purchased from ATCC (Manassas, VA, USA). A2780 was purchased from ECACC (Porton Down, Wiltshire, UK). Cells were confirmed to be pathogen-free (Mycoplasma testing, Lonza, L108-318). MCF-7 cells were maintained in DMEM (Thermo Fisher Scientific, Waltham, MA, USA, Gibco™, 31053028) with 10% FBS (Invitrogen, Carlsbad, CA, USA, 10091148), GlutaMAX (Thermo Fisher Scientific, Gibco™, 35050061) and Penicillin-Streptomycin (Welgene, Gyeongsansi, South Korea, LS202-02), and for short-term treatment, DMEM (Gibco™, 31053028) with 10% charcoal stripped FBS (Gibco™, A3382101), and GlutaMAX (Gibco™, 35050061) was used. HCT116 were maintained in McCoy’s 5A medium (ATCC, 30-2007) with 10% FBS (Invitrogen, 10091148), DMS53 were maintained in Waymouth’s MB 752/1 medium (Merck, Darmstadt, Germany, W1625) with 10% FBS (Invitrogen, 10091148), and H82, H292, and A2780 were maintained in RPMI 1640 (ATCC, 30-2001) with 10% FBS (Invitrogen, 10091148). Cells were cultured at 37, 5% CO2. All animal experiments were performed in compliance with protocols approved by the Institutional Animal Care and Use Committee (IACUC) of WuXi AppTec and complied with the guidelines by the Association for Assessment and Accreditation of Laboratory Animal Care (AAALAC). Q901 and Bio-QS (Bio-QS1169; a biotinylated Q901 analog) were synthesized by Qurient (Seongnam-si, South Korea). Bortezomib (BTZ) and Topotecan (TPT) were purchased from Sigma-Aldrich (St. Louis, MO, USA, 5.04314, and T2705). Trastuzumab deruxtecan (T-DXd) and sacituzumab govitecan (SG) were purchased from Evidentic GmbH (Potsdam, Germany).

The following antibodies were used for immunoblotting: total CDK7 (Cell Signaling Technology, Danvers, MA, USA, 2090S, RRID: AB_2077140), MYC (Cell Signaling Technology, 13987, RRID: AB_2631168), E2F1 (Cell Signaling Technology, 3742, RRID: AB_2096936), MYC S62P (Cell Signaling Technology, 13748, RRID: AB_2687518), MYC T58P (Cell Signaling Technology, 46650), Rb S795P (Cell Signaling Technology, 9301, RRID: AB_330013), Rb S780P (Cell Signaling Technology, 9307, RRID: AB_330015), MAX (Cell Signaling Technology, 4739, RRID: AB_2281777), T-H2AX (Santa Cruz, Dallas, TX, USA, sc-517336, RRID: AB_3675923), P-H2AX (Cell Signaling Technology, 2577, RRID: AB_2118010), Ubiquitin (Santa Cruz, sc-8017, RRID: AB_628423), TOP1 (BD Biosciences, San Jose, CA, USA, 556597, RRID: AB_396474), β-actin (Santa Cruz, sc-47778, RRID: AB_626632), Affinity Purified Goat Anti-Rabbit IgG (H+L)-HRP (Bio-Rad, Hercules, CA, USA, 1706515, RRID: AB_11125142), and Affinity Purified Goat Anti-Mouse IgG (H+L)-HRP (Bio-Rad, 1706516, RRID: AB_11125547).

The following antibodies were used for ChIP-seq: pan pol II (Cell Signaling Technology, 2629 RRID: AB_2167468), Ser5P (Abcam, Cambridge, UK, ab5408, RRID:AB_304868), Ser2P (Abcam, ab5095, RRID:AB_304749), CDK7 (Bethyl, Montgomery, TX, USA, A300-405A, RRID: AB_2275973), MYC (Cell Signaling Technology, 13987, RRID: AB_2631168), E2F1 (Cell Signaling Technology, 3742, RRID: AB_2096936), and TOP1 (Abcam, ab109374, RRID: AB_10861978).

### Inhibitor treatment

To assess the durability of CDK7 occupancy after Q901 removal, cells were treated with 6 nM Q901 for 4 h, washed with fresh medium, and incubated for 4, 24, or 48 h. For the short-term condition, MCF-7 cells were treated with DMSO or 100 nM Q901 for 1 h followed by 1 h of EtOH/E2 (estrogen) treatment (the E2 condition was not used in this study). For long-term treatment, MCF-7 cells were treated with DMSO or 100 nM Q901 for 4, 24, 48 or 72 h. For TOP1 ChIP-seq, cells were treated with 100 nM Q901 for 2 h followed by 1 μM TPT for 6 h. For DUST assay, cells were treated with 1 μM Q901 for 24 h followed by 1 μM TPT for 0.5, 1, or 2 h. 1 μM BTZ was treated concurrently with Q901. For the detection of TOP1 protein from whole cell lysates, MCF-7 cells were pre-treated with DMSO or Q901 100 nM for 2 h, followed by treatment with TPT 1 μM for 0, 1, 6, or 24 h.

### NMR Analysis

¹HNMR spectrum was recorded at 298 K (VT-NMR at 353 K) using a Bruker Avance III™ HD 400 MHz NMR spectrometer equipped with a broadband observe (BBO) probe (Bruker BioSpin GmbH, Rheinstetten, Germany). Chemical shifts (δ) are reported in parts per million (ppm), calibrated to residual non-deuterated solvent signals. Multiplicities are abbreviated as s (singlet), d (doublet), t (triplet), q (quartet), m (multiplet), and br (broad). Coupling constants (J) are reported in hertz (Hz), and integrals are provided for ¹HNMR spectra.

### 410 Kinases inhibition profile and 28 CDKs titration

A radiometric protein kinase assay service was provided by Reaction Biology (Freiburg, Germany) to measure the kinase activity of 410 protein kinases and 28 CDKs.

In the kinase inhibition assay, the residual activity for each compound was calculated using the following formula: Residual Activity (%) = 100 X [(signal of compound – low control) / (high control – low control)]. The selectivity score of the compound at the tested concentrations was calculated for a residual activity < 50 %, that is, inhibition of > 50 %. The selectivity score for a particular compound at a particular concentration was calculated using the following formula: Selectivity Score = (count of data points < 50 %) / (total number of data points).

In the 28 CDK titration assay, the residual activity and IC50 values of the compound were calculated using Quattro Workflow V3.1.1 (Quattro Research GmbH, Munich, Germany). The ATP concentration used in each kinase reaction was set to the apparent ATP-Km value. Dose-response curve fitting was performed using a four-parameter logistic regression model in GraphPad Prism 10.4.1 (GraphPad Software, Boston, MA, USA).

### Q901 covalent binding site evaluation

The recombinant CAK complex (CDK7/Cyclin H/MAT1, Thermo Fisher Scientific, PV3868) was incubated with Q901 or DMSO for 1 h at room temperature, and the reaction complexes were resolved by SDS-PAGE. The bands corresponding to CDK7 were excised for in-gel digestion. Denaturation and reduction were performed using TCEP heated at 55°C, followed by ArgC digestion (1:50 ArgC:Protein ratio, at 37°C for 18 h) and trypsin digestion (1:50 ArgC:Protein ratio and 1:5 Trypsin:Protein ratio, at 37°C for 18 h). The extracted peptides were loaded and eluted using an HPLC system, and then separated through a linear gradient (0% to 90% mobile phase B in 30 min) at a flow rate of 0.5 mL/min using an Agilent PoroShell 120 EC-C18 column (2.1 mm × 50 mm) at 60°C and introduced into a QTRAP® 5500 LC-MS/MS System (SCIEX, Redwood city, CA, USA) in positive ion mode and run in data-dependent acquisition (DDA) mode, after which the four most abundant precursors were subjected to MS/MS analysis. Data acquisition was performed over an m/z range of 100–1,200, and peak integrations for doubly, triply, and quadruply charged precursors of the doubly carbamidomethylated Cys312-containing peptide were analyzed using Skyline software 24.1 (Seattle, WA, USA). The carbamidomethylated Cys304-containing CDK7 peptides, RPGPTPGCQLP (residue 298-308) was used for normalization.

### CDK7 target occupancy assay

A2780 cells were seeded into 10 cm dishes and treated with Q901 or DMSO as the indicated concentrations for 4 h. Cells were washed twice with cold PBS, lysed with 200 μL RIPA buffer (Sigma-Aldrich, R0278), and incubated on ice for 30 min. Lysates were scrapped, centrifugated at 12,000 rpm for 10 min at 4°C, and protein concentration was determined using a BCA assay. A2780 cell lysates were incubated with 1 μM of Bio-QS overnight at 4 with gentle rocking, then incubated for 3 h at room temperature (RT) to enhance covalent binding. Bio-QS binds strongly to CDK7 and has the same covalent cassette as Q901 does. After incubation, 200 μL of 50% streptavidin agarose bead slurry (Thermo Fisher Scientific, 20353) was added to the cell lysates, and the samples were incubated overnight at 4°C with gentle rocking for immunoprecipitated (IP). The following day, the samples were pelleted by microcentrifugation (14,000 g, 30 sec, 4°C), washed three times with ice-cold RIPA buffer, and the supernatant was collected prior to washing. The pellet was resuspended in 2X SDS sample buffer, vortexed, microcentrifuged for 30 sec, heated at 95-100°C for 5 min, and centrifuged again at 14,000 g for 1 min. IP reaction samples were loaded on 4-12% Bis-Tris gels as follows: equal volume of IP-reacted samples, input protein, and Bio-QS-incubated lysate for total CDK7 detection. Proteins were transferred to PVDF membranes, blocked with 5% BSA, and immunoblotted with a CDK7 primary antibody (Cell Signaling Technology, 2090S) followed by a secondary HRP-conjugated antibody. Signals were detected using ECL and analyzed using Image Studio Lite Ver5.2 (LI-COR Biotechnology, Lincoln, NE, USA). Quantitative data of immunoblotting bands were generated by GraphPad Prism 10.4.1.

### CellTiter-Glo assay (CTG assay)

Cells were seeded into 96-well plates and treated with compounds at the indicated concentrations and time points. After incubation, cell viability was measured using either CellTiter-Glo luminescent cell viability assay (Promega, Madison, WI, USA, G7572) according to the manufacturer’s instructions. Plates were equilibrated to RT for 30 min before adding 50 μL of CellTiter-Glo reagent to each well. Luminescence was measured using EnVision Multi Label Reader (Perkin Elmer, Waltham, MA, USA). Dose-response curves were fitted using a nonlinear regression model with a sigmoidal dose-response function in GraphPad Prism 10.4.1. The absolute IC□□ values were calculated using the dose-response curves generated in XL-fit 5 (IDBS Software, Woking, UK) and fitted with the four-parameter logistic equation: fit = A + ((B - A) / (1 + ((C / x) ^ D))) where A and B represent the bottom and top plateaus of the curve, respectively; C is the IC□□ value, x is the compound concentration, and D is the Hill slope.

### Immunoblotting assay

Cell lysates were obtained using RIPA buffer (BIOSESANG, Yongin-si, South Korea, R2002) containing protease inhibitor (GenDEPOT, Baker, TX, USA, P3100) and phosphatase inhibitor cocktails (GenDEPOT, P3200). Samples were collected and kept on ice for 30 min. After centrifugation at 12,000 rpm for 10 min at 4°C, the supernatant was collected for further analysis. Protein concentrations were determined using a Pierce™ BCA Protein Assay Kit - Reducing Agent Compatible (Thermo Fisher Scientific, Pierce, 23250). Equal amounts of protein were loaded onto SDS-PAGE gels and electrophoresed at 100 V for 1 h 45 min. Proteins were transferred onto PVDF membranes (MilliporeSigma, Burlington, MA, USA) and blocked with 5% BSA in TBST for 1 h at RT. Membranes were incubated with primary antibodies overnight at 4°C, washed 3 times for 10 min each on a shaker, and incubated with HRP-conjugated secondary antibodies for 45 min at RT. The membranes were washed again and soaked in substrate working solution (Merck, WBKLS0100) for 10 sec. Chemiluminescent signals were detected using Image Quant LAS 4000 (PerkinElmer, Waltham, MA, USA). Quantitative data of immunoblotting bands were generated with ImageJ (1.54) and GraphPad Prism 10.4.1.

### ChIP (Chromatin Immunoprecipitation)-seq

ChIP-seq was performed as previously described with minor changes (*45*). After Q901 or TPT was treated, MCF-7 cells were crosslinked for 10 (MYC, E2F1 and TOP1) or 15 (other proteins) min at RT by crosslinking buffer (50 mM HEPES pH 8.0, 100 mM NaCl, 1 mM EDTA pH 8.0, 0.5 mM EGTA pH 8.0) with 1% formaldehyde (Sigma-Aldrich, F8775-4X25ML) followed by quenching with 125 mM glycine (Daejung, Siheung-si, South Korea, 4068-4105) for 5 min. Cells were washed with ice cold 1X PBS and pellets were stored at -80. Frozen pellets were lysed with buffer I (50 mM HEPES pH 7.5, 140 mM NaCl, 1 mM EDTA pH 8.0, 10% glycerol, 0.5% NP-40, 1X protease inhibitor (GenDEPOT, P3100), and 1X phosphatase inhibitor (GenDEPOT, P3200)). The pellets were resuspended in modified buffer II/III (10 mM Tris-HCl pH 8.0, 300 mM NaCl, 0.1% DOC (sodium-deoxycholate), 1% Triton X-100, 1 mM EDTA pH 8.0, 1 mM EGTA pH 8.0, 1X protease inhibitor, and 1X phosphatase inhibitor) and sonicated at 4. Sonicated cells were incubated with antibody overnight at 4 with rotation. Agarose beads were washed 3 times with modified buffer II/III. Washed beads were added to sonicated cells and incubated for 2 h at 4 with rotation. Bead-antibody-conjugated cells were washed 2 times with ice-cold low salt washing buffer (20 mM Tris-HCl pH 8.1, 0.1% SDS, 1% Triton X-100, 2 mM EDTA pH 8.0, 150 mM NaCl), 2 times with ice-cold high salt washing buffer (20 mM Tris-HCl pH 8.1, 0.1% SDS, 1% Triton X-100, 2 mM EDTA pH 8.0, 500 mM NaCl), 2 times with ice-cold LiCl washing buffer (10 mM Tris-HCl pH 8.1, 250 mM LiCl, 1% NP-40, 1 mM EDTA pH 8.0, 1% DOC), and 1 time with RT 1X TE buffer. DNA was eluted with elution buffer (10 mM Tris-HCl pH 8.0, 1 mM EDTA pH 8.0, 1% SDS) and reverse-crosslinked for 5 h at 65. RNA and protein were removed using RNase A (BIOSESANG, R1007) and Proteinase K (NEB, Ipswich, MA, USA P8107S), respectively and DNA was purified with UltraPure™ Phenol:Chloroform:IsoamylAlcohol (25:24:1, v/v) (Thermo Fisher Scientific, 15593031) and MinElute PCR purification kit (QIAGEN, 28006).

ChIP-seq libraries were generated with NEBNext End Repair Module (NEB, E6050L), NEBNext dA-Tailing Module (NEB, E6053L), Quick Ligation Kit (NEB, M2200L), NEBNext Multiplex Oligos for Illumina (NEB, E7335S, E7500S, E7710S), and Phusion™ High-Fidelity DNA Polymerase (NEB, M0530L) according to the manufacturer’s protocol. ChIP-seq samples were sequenced on Illumina NextSeq500 (Illumina, San Diego, CA, USA) with 75bp single-end reads.

### mRNA-seq

Total RNA was prepared from MCF-7 cells after Q901 treatment using TRIzol (Invitrogen, 15596026) according to the manufacturer’s protocol. mRNA-seq library was constructed using the MGIEasy RNA Directional Library Prep Kit (MGI, Shenzhen, China) according to the manufacturer’s instructions and sequenced on MGISEQ-2000 (MGI) with 100bp paired-end reads.

### fastGRO (fast Global Run-On)

fastGRO sample preparation was performed as previously described with minor modifications (*46*). After Q901 and TPT were treated, nuclei were isolated with ice cold swelling buffer (10 mM Tris-HCl pH 7.5, 2 mM MgCl_2_, 3 mM CaCl_2_, 2 U/ml SUPERase-In RNase inhibitor (Invitrogen, AM2696)), swelling buffer with glycerol (10 mM Tris-HCl pH 7.5, 2 mM MgCl_2_, 3 mM CaCl_2_, 10% glycerol, 2 U/ml SUPERase-In RNase inhibitor), and lysis buffer (10 mM Tris-HCl pH 7.5, 2 mM MgCl_2_, 3 mM CaCl_2_, 10% glycerol, 1% NP-40, 2 U/ml SUPERase-In RNase inhibitor). Nuclei were stored in freeze buffer (50 mM Tri-HCl pH 8.0, 40% glycerol, 5 mM MgCl_2_, 0.1 mM EDTA pH 8.0, 2 U/ml SUPERase-In RNase inhibitor) at -80. At run-on step, nuclei were incubated with nuclear run-on buffer including 4-thio-UTP (10 mM Tris-HCl pH 8.0, 5 mM MgCl_2_, 1 mM DTT, 300 mM KCl, 500 uM ATP, 500 uM GTP, 2 uM CTP, 500 uM 4-S-UTP, 1% sarkosyl, SUPERase-In RNase inhibitor) for 7 min. After total RNA precipitation, RNA was fragmented by sonication and biotinylated using EZ-link HPDP-Biotin (Thermo Fisher Scientific, A35390). Biotinylated RNA was extracted using RNA precipitation and streptavidin-coupled magnetic beads (Dynabeads MyOne Streptavidin C1, Invitrogen, 65001).

Library preparation was performed following the qPRO-seq protocol (*47*). Following the ligation of the 3’ adapter, samples were treated with RppH (NEB, M0356S) and PNK (NEB, M0201S), and 5’ adapter was ligated. RNA was purified using streptavidin-coupled magnetic beads and washed between incubation steps with high-salt buffer (50 mM Tris-HCl pH 7.4, 2 M NaCl, 0.5% Triton X-100, 1 mM EDTA pH 8.0, 40 U/10 mL SUPERase-In RNase inhibitor) and low-salt buffer (50 mM Tris-HCl pH 7.4, 0.1% Triton X-100, 1 mM EDTA pH 8.0, 40 U/10 mL SUPERase-In RNase inhibitor). The biotinylated RNA was converted to DNA with reverse-transcription step (Maxima H-reverse transcriptase, Thermo Fisher Scientific, EP0751) and amplified (Q5 High-Fidelity DNA Polymerase, NEB, M0491L) for sequencing. fastGRO samples were sequenced on NovaSeq6000 (Illumina) with 100bp paired-end reads.

### DUST (Detection of Ubiquitylated and SUMOylated TOP1 DNA-Protein Crosslinks) assay

The DUST assay was performed according to the previously described procedure (*25, 48*) with slight modifications using MCF-7 cells treated with TPT or a combination of TPT and Q901. Cells were washed with PBS, lysed with DNAzol (Invitrogen, 10503027) containing 1X protease inhibitor (GenDEPOT, P3100) and 1mM DTT, and nucleic acids were precipitated with ethanol. The precipitated nucleic acids were resuspended in water, heated at 37°C for 30 min, and sonicated. After centrifugation at 20,000 g for 5 min, the supernatant was collected for spectrophotometric quantification. Each sample was digested with 1000 units of micrococcal nuclease (NEB, M0247), followed by SDS-PAGE. Detection of ubiquitylated TOP1-DPCs was carried out using an ubiquitin antibody.

### T-DXd Serum Stability Assay

The serum stability of trastuzumab-DXd (T-DXd) was evaluated in male CD-1 mouse serum (Vital River Laboratory Animal Technology Co.,Ltd, Beijing, China). Blood was collected via cardiac puncture under anesthesia, and serum was isolated by centrifugation (3000 × g, 10 min, 4°C), aliquoted, and stored at −80°C until use. T-DXd was spiked into the mouse serum at a final concentration of 100 µg/mL and incubated at 37°C for 0, 1, 4, 7, 14, and 21 days. Buffer controls were prepared by incubating 100 µg/mL of T-DXd in PBS (Gibco, 2661454) containing 1% Bovine Serum Albumin (BSA; BBI J711BA007). High-quality control (HQC) samples were prepared by spiking T-DXd into mouse serum at the same concentration. Samples were processed using High Capacity Magne™ Streptavidin Beads (Promega, 576739), charged with Goat Anti-human Biotinylated IgG (Promega, 571395). Serum samples (20 µL) were diluted with 130 µL of bind/wash buffer (200 mM NaCl in PBS, prepared with NaCl from Bidepharm, EKZ867, and PBS from Gibco, 2661454), and incubated with the beads at 1100 rpm for 120 min at room temperature. Beads were washed twice with bind/wash buffer, once with PBS, and once with purified water (ELGA, NA). For elution and reduction, 50 µL of elution buffer (0.1% trifluoroacetic acid in water; J&K, L910X51) was added, and the mixture was shaken at 1100 rpm for 30 min at room temperature. The eluate was neutralized with 5 µL of neutralization buffer (1 M ammonium bicarbonate; LingFeng, 20231115) and 5 µL of DTT (TCI, OJWPT-DA). Samples were shaken for another 30 min at room temperature and centrifuged at 3220 g for 5 min. LC-MS/MS analysis was performed using a Waters VION QTof mass spectrometer operated in sensitivity mode (ESI+, m/z 800–2500). The source temperature was set to 150°C, with a desolvation temperature of 600°C, desolvation gas flow at 700 L/h and cone gas flow at 50 L/h. The capillary voltage was 2.75 kV, the sample cone voltage was 25 V, and the source offset voltage was 40 V. LockSpray was applied using m/z 785.8421 as the reference. Chromatographic separation was performed on an Agilent PLRP-S column (1 × 50 mm, 5 µm, 1000 Å; Agilent Technologies, AT-PL1312-1502) at 60°C with a flow rate of 0.3 mL/min. The mobile phases consisted of 0.1% formic acid (Thermo Fisher, A117) in water (A) and 0.1% formic acid in acetonitrile (Merck, 1.000.30.4008) (B). The gradient elution was programmed from 10% to 85% B over 8 min, followed by re-equilibration to initial conditions by 12 min. The Drug-to-Antibody Ratio (DAR) values of T-DXd over time were measured. This study was performed in duplicate. This study was performed in duplicate. Data represent mean ± standard deviation (SD).

### Cell line-derived xenograft (CDX) models

BALB/c nude mice were purchased from Shanghai SIPPR/BK Laboratory Animal Co., LTD. (Shanghai, China). For in vivo efficacy studies, HCT116 cells or H292 cells were resuspended in PBS and Matrigel (1:1) and implanted subcutaneously. When tumors reached an average size of 117 mm³ (HCT116) or 140 mm³ (H292), mice were randomized into four groups (n = 8 per group) and treated with Q901 (3 mg/kg or 10 mg/kg, intraperitoneally once daily), T-DXd (10 mg/kg, single intravenous administration on day 0), SG (3 mg/kg, single intravenous administration on day 1 and 8), or a combination thereof. Tumor volumes were measured twice a week using calipers, and the tumor growth inhibition (TGI) was calculated using the following formula: Tumor volumes were measured twice weekly after randomization in two dimensions, using a caliper. TGI was calculated for each group using the formula: TGI (%) = [1-(Ti-T0)/ (Vi-V0)] ×100; Ti is the average tumor volume of the treatment group on the given day, T0 is the average tumor volume of the treatment group on the day when the treatment was started, Vi is the average tumor volume of the vehicle control group on the same day with Ti, and V0 is the average tumor volume of the vehicle group on the day of treatment start. Statistical significance was calculated using two-way ANOVA followed by Tukey’s multiple comparison test in GraphPad Prism 10.4.1.

### SynergyFinder™

SynergyFinder™ was performed by Oncolines® (Oss, Netherlands). Cells were treated with Q901, T-DXd, or SG for 72 h, and cell viability was assessed to determine IC50 of Q901, T-DXd, or SG in the respective cell lines. Since HCT116 cells were insensitive to T-DXd, 10 µg/mL of T-DXd was mixed with various concentrations of Q901 in HCT116 cells. In H292 cells, which are sensitive to SG, stock solutions of Q901 and SG were diluted to their respective IC50 concentrations, as determined in single-agent experiments, and then mixed at a 1:4 ratio (Q901:SG = 1:4). The mixtures, Q901, and SG were further diluted to generate a 7-point dose-response series in duplicate. For single agents, the final assay concentration range is from 10 to 0.01 times their IC50 (10 and 0.01 equivalents). Dose-response curves were fitted using nonlinear regression with a sigmoidal dose-response model in GraphPad Prism 10.4.1 or XL-fit 5. The absolute IC□□ values were determined from the dose-response curves generated in XL-fit 5 using the four-parameter logistic equation: fit = A + ((B - A) / (1 + ((C / x) ^ D))) where A and B represent the bottom and top plateaus of the curve, C is the IC□□ value, x is the compound concentration, and D is the Hill slope.

### Data analysis

#### ChIP-seq analysis

ChIP-seq data were trimmed with Trim Galore (0.6.7) (https://github.com/FelixKrueger/TrimGalore) and aligned to hg38 (v36) using Bowtie2 (2.4.5) (https://github.com/BenLangmead/bowtie2). The bigWig files were generated by HOMER (v4.11) (http://homer.ucsd.edu/homer/) makeUCSCfile and visualized with UCSC genome browser. For pan RNAPII, Ser5P, and Ser2P ChIP-seq, read counting was generated by HOMER analyzeRepeats.pl and DESeq2 (1.40.0) (https://bioconductor.org/packages/DESeq2/). For CDK7, MYC, E2F1, and TOP1 ChIP-seq, peakcalling was performed with MACS3 (3.0.0b1) (https://github.com/macs3-project/MACS) and IDR (2.0.3) (https://github.com/hbctraining/Intro-to-ChIPseq).

#### mRNA-seq analysis

mRNA-seq data were trimmed with Trim Galore and aligned to hg38 (v36) using STAR (2.7.8a_2021-03-08) (https://github.com/alexdobin/STAR). The bigWig files were generated by deepTools bamCoverage (3.5.1) and visualized with UCSC genome browser. The expression level was calculated with RSEM rsem-calculate-expression (v1.3.1) (https://github.com/deweylab/RSEM), and DESeq2.

#### fastGRO analysis

fastGRO data were trimmed with fastp (0.23.4) (https://github.com/OpenGene/fastp) and rRNA was removed. The trimmed reads were aligned to hg38 using Bowtie2 and deduplicated with UMI-tools (1.1.4) (https://github.com/CGATOxford/UMI-tools). The bigWig files were generated by deepTools (https://github.com/deeptools/deepTools) bamCoverage (3.5.1) and visualized with UCSC genome browser.

#### Average plot and heatmap

Average plot and heatmap were generated by deepTools computeMatrix, and plotHeatmap (3.5.1).

#### Pathway analysis

GSEA (Gene Set Enrichment Analysis) was performed using R package fgsea (1.26.0) (https://github.com/alserglab/fgsea). The datasets were used from MSigDB (v2023.1) (https://www.gsea-msigdb.org/gsea/msigdb). GO analysis of enhancer target genes and hallmark pathway enrichment analysis of intersecting genes from E2F1 and MYC cluster1 was performed using the Metascape (https://metascape.org/).

#### Enhancer calling

The enhancers were defined using H3K27ac ChIP-seq from GSE62229 (*32*). ChIP-seq data were processed same as CDK7, MYC, E2F1, and TOP1 ChIP-seq except for peakcalling step (MACS3 option: --broad). ROSE (https://github.com/stjude/ROSE) was used to define SEs with H3K27ac peaks (-s 12500 –t 2000). The defined enhancers that did not overlap with SE were defined as TEs. CDK7-bound enhancers were defined as enhancers that overlapped with peaks showing increased CDK7 binding upon Q901 treatment compared to DMSO. CDK7-nonbound enhancers were defined as the remaining enhancers after excluding CDK7-bound enhancers from the total enhancer set. The target protein-coding genes of SE and TE were defined as protein-coding genes located within a 50 kb region from the start or end of SE/TE. To ensure that the four groups did not overlap, priority was given to SE CDK7-bound. For target genes containing at least one SE, those with at least one CDK7-bound SE were classified as SE CDK7-bound target genes, while those without any CDK7-bound SE were classified as SE CDK7-nonbound target genes. The same classification criteria were applied to TE target genes that did not contain any SE.

### Statistical Analysis

Statistical analyses were performed using GraphPad Prism 10.4.1. Statistical approaches are specified for each experiment and only p-value ≤ 0.05 are considered significant.

## Supporting information

supplemental figurea and table

## Acknowledgments

We thank Won-Gyun Ahn for designing the T-DXd serum stability assay.

## Funding

This work was supported by the National Research Foundation of Korea (NRF) grant funded by the Korean government (MSIT) (RS-2023-00265581 to T.-K.K.), (RS-2023-NR077236 to T.-K.K.). This study was also supported by the Samsung Science & Technology Foundation (SSTF-BA2102-09 to T.-K.K.), the Multitasking Macrophage Research Center (RS-2023-00217798 to T.-K.K.), and the Korea Basic Science Institute (National Research Facilities and Equipment Center) grant funded by the Ministry of Education (RS-2021-NF000572 to T.-K.K.). Y.S. received research support from the National Cancer Institute (4R00CA273171-02). A.T. was supported by the intramural programs of the Center for Cancer Research, National Cancer Institute (ZIA BC 011793), reports research funding to the institution from the following entities: EMD Serono Research & Development Institute; AstraZeneca; Tarveda Therapeutics; Immunomedics and Prolynx.

## Author contributions

Conceptualization: H.J., Y.L., D.Y., Y.J., H.K., M.S., D.P., S.J.L, J.S.K., K.N., and T.-K.K.

Data curation: H.J., Y.L., D.Y., Y.J., H.K., D.U., M.S., D.P., S.J.L, and J.S.K.

Formal Analysis: H.J., Y.L., D.Y., Y.J., H.K., D.U., and S.J.L.

Funding acquisition: K.N., and T.-K.K.

Investigation: H.J., Y.L., D.Y., Y.J., H.K., M.S., D.P., and J.S.K.

Methodology: L.E., Y.S., and A.T.

Project administration: H.J., Y.L., D.Y., J.J.K., S.J.L, K.N., and T.-K.K.

Resources: K.N., and T.-K.K.

Supervision: K.N., and T.-K.K.

Validation: H.J., Y.L., D.Y., S.J.L., K.N., and T.-K.K.

Visualization: H.J., and Y.L.

Writing – original draft: H.J., and Y.L.

Writing – review & editing: H.J., Y.L., S.J.L, K.N., and T.-K.K.

## Competing interests

Y.L., D.Y., Y.J., H.K., M.S., D.P., J.J.K., S.J.L, J.S.K., and K.N. are employees and hold stock in Qurient Co. Ltd., which develops Q901 related to the research discussed in this paper. T.-K.K received research funding from Qurients Co., Ltd. The remaining authors declare no competing interests.

## Data and materials availability

The ChIP-seq (Pan RNAPII, Ser5P, Ser2P, CDK7, MYC, E2F1, and TOP1), mRNA-seq, and fastGRO data generated in this study have been deposited in the NCBI Gene Expression Omnibus (GEO) database under the accession codes GSE292907, GSE292860, and GSE292846. H3K27ac ChIP-seq data used in this study are available in GEO under the accession code GSE62229.

## Supplementary Materials

**Fig. S1.**
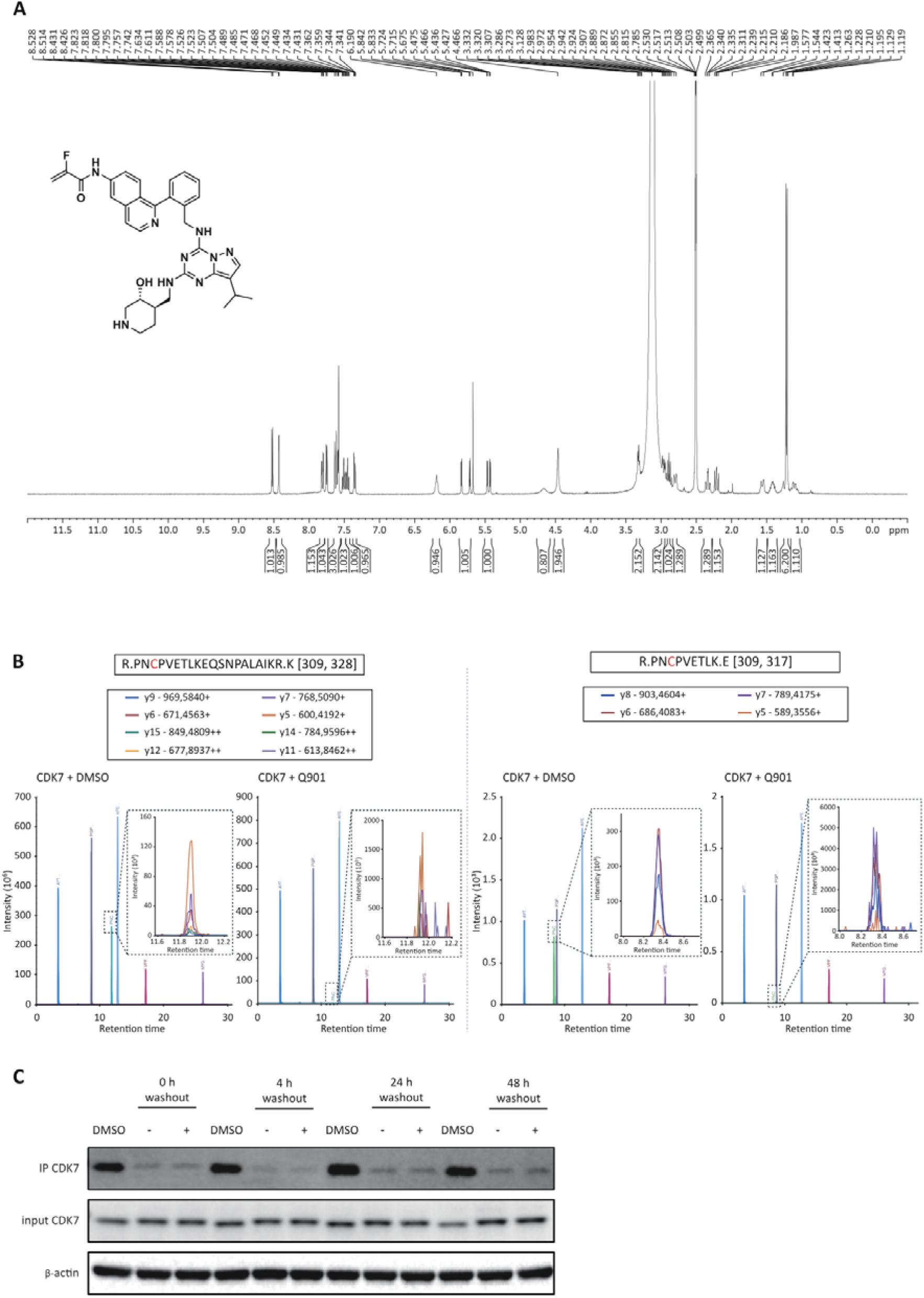
Biochemical and proteomic validation of Q901 specificity. **A**) ^1^H NMR spectrum of Q901 was acquired using variable temperature (VT) NMR in DMSO-d□. (**B**) Targeted proteomics analysis to determine the Q901 binding sites on CDK7. The recombinant CAK trimeric complex was incubated with Q901 or DMSO, followed by protease digestion and peptide mapping via LC-MS/MS. Chromatograms show peptide fragments generated by ArgC (Clostripain) digestion (right) and ArgC/Trypsin digestion (left). The expanded boxes highlight the peak of C312 containing peptides, which are reduced following Q901 treatment, indicating covalent modification at this site. (**C**) Representative Western blot images from the pulse-chase assay described in Fig. 1F. A2780 cells were treated with 6 nM Q901 for 4 h and then divided into two groups. One group (- wash out) remained in the Q901-containing medium for continuous incubation, while the other group (+ wash out) underwent a drug washout, where the medium was completely removed and replaced with fresh drug-free medium before further incubation for the indicated times. Bio-QS-labeled CDK7 was immunoprecipitated using streptavidin agarose beads (SA), and the levels of free CDK7 were analyzed by immunoblotting. These images in this figure were quantified in Fig. 1F.

**Fig. S2.**
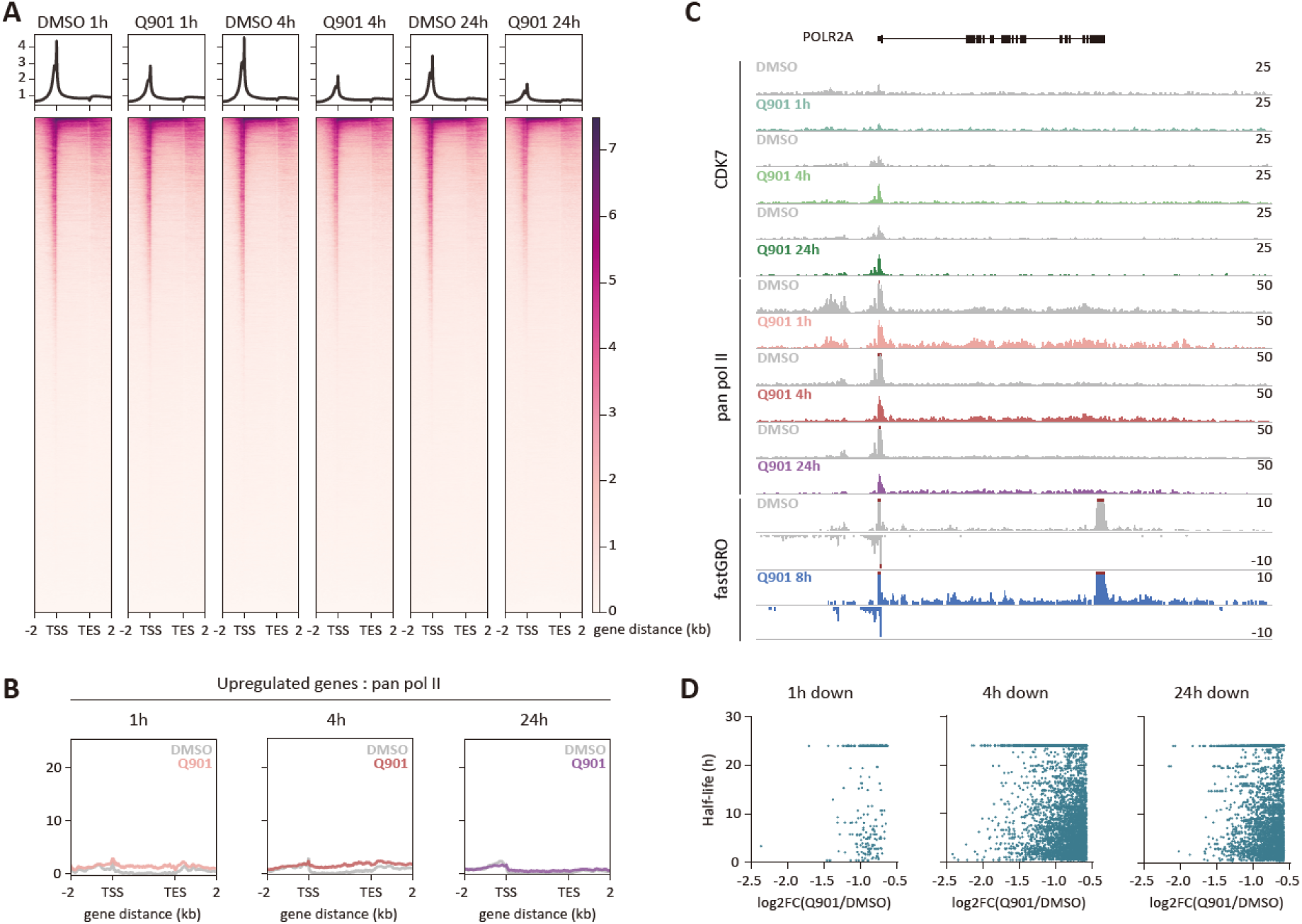
Global and gene-specific changes in RNAPII occupancy following Q901 treatment. (**A**) Heatmap of pan RNAPII ChIP-seq for all annotated genes (n = 2, p-value ≤ 0.05). (**B**) Average ChIP-seq plots of pan RNAPII for upregulated genes across various time points following Q901 treatment, presented on the same scale as those for downregulated genes in Fig. 2D. (**C**) Track image of POLR2A, an example of an upregulated gene by Q901 treatment. (**D**) Scatter plots showing the correlation between the mRNA half-lives of downregulated genes and log2FC values of pan RNAPII ChIP-seq for the corresponding genes.

**Fig. S3.**
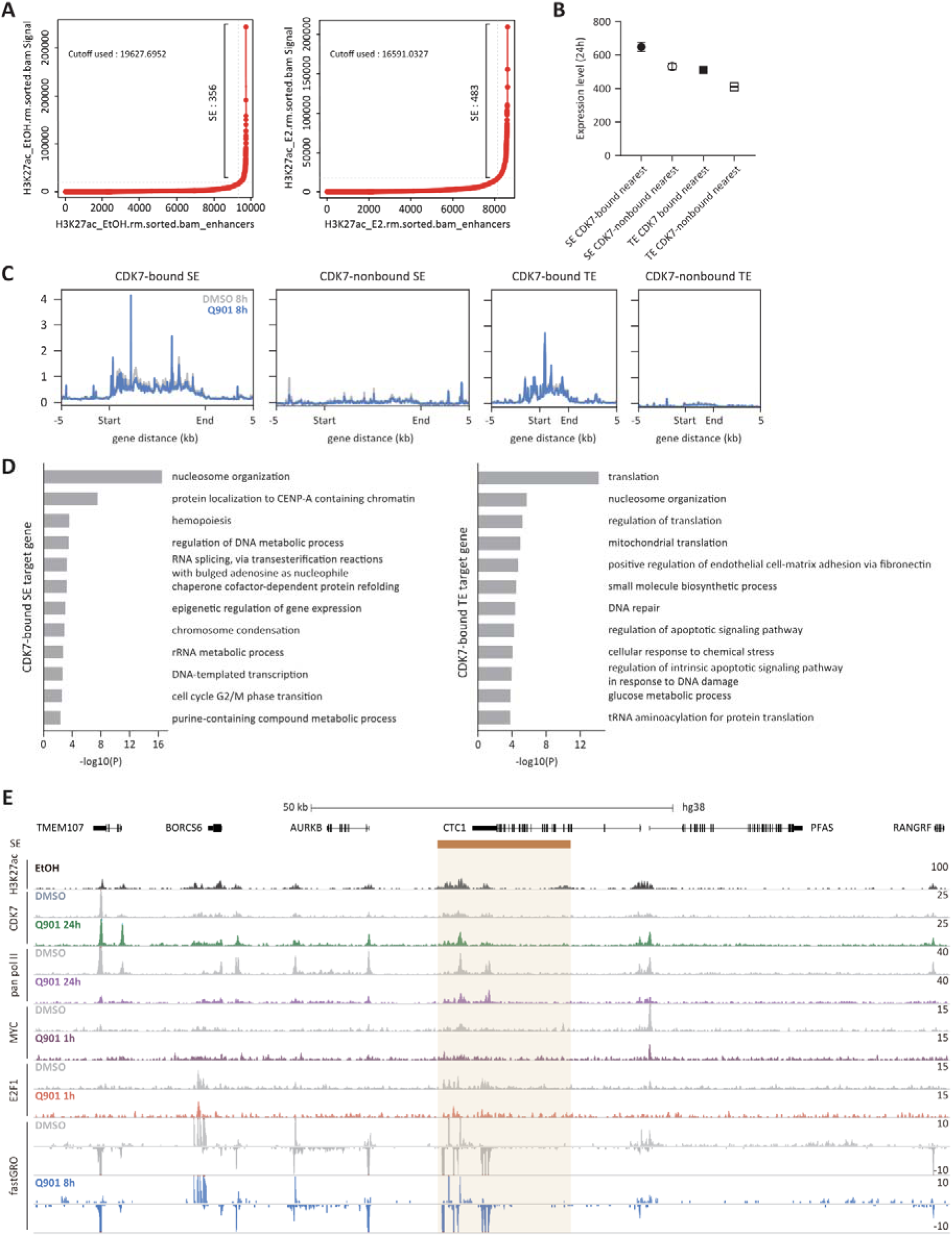
Transcription profiles of enhancer target genes. (**A**) The results of SE calling using the ROSE program with H3K27ac ChIP-seq (GSE62229). (**B**) Expression levels of enhancer target genes (pan RNAPII ChIP-seq; n = 2, Q901 1h treatment condition, data represent mean ± SEM). (**C**) Average fastGRO signals of four enhancer groups. (**D**) GO analysis results of target genes regulated by CDK7-bound SE and CDK7-bound TE. (**E**) Track images showing ChIP-seq signals for H3K27ac, CDK7, pan RNAPII, MYC, and E2F1, along with fastGRO, at a representative CDK7-bound SE region (highlighted in yellow) and it associated target genes.

**Fig. S4.**
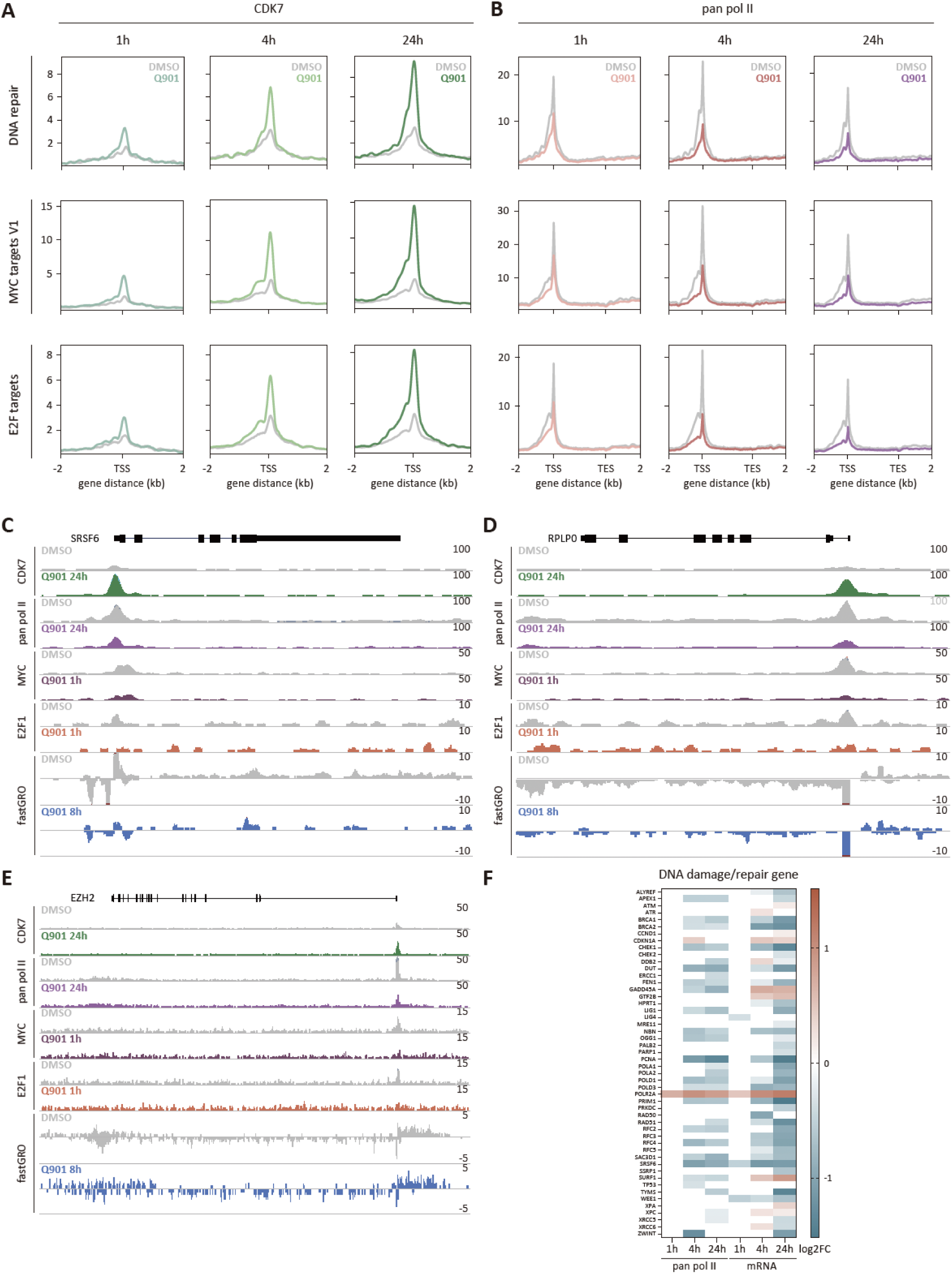
CDK7 and RNAPII ChIP-seq signal at DNA Repair, MYC, and E2F target genes. (**A** and **B**) Average ChIP-seq signal plots of CDK7 (A) and pan RNAPII (B) for gene sets related to DNA repair, MYC targets V1, and E2F targets. (**C**) Track image of SRSF6 gene, a representative gene from the DNA repair pathway. (**D**) Track image of RPLP0 gene, a representative gene from the MYC targets V1 pathway. (**E**) Track image of EZH2 gene, a representative gene from the E2F targets pathway. (**F**) Heatmap showing log2FC values of DNA damage/repair genes expression from pan RNAPII ChIP-seq and mRNA-seq data (right; n = 3, FDR ≤ 0.1).

**Fig. S5.**
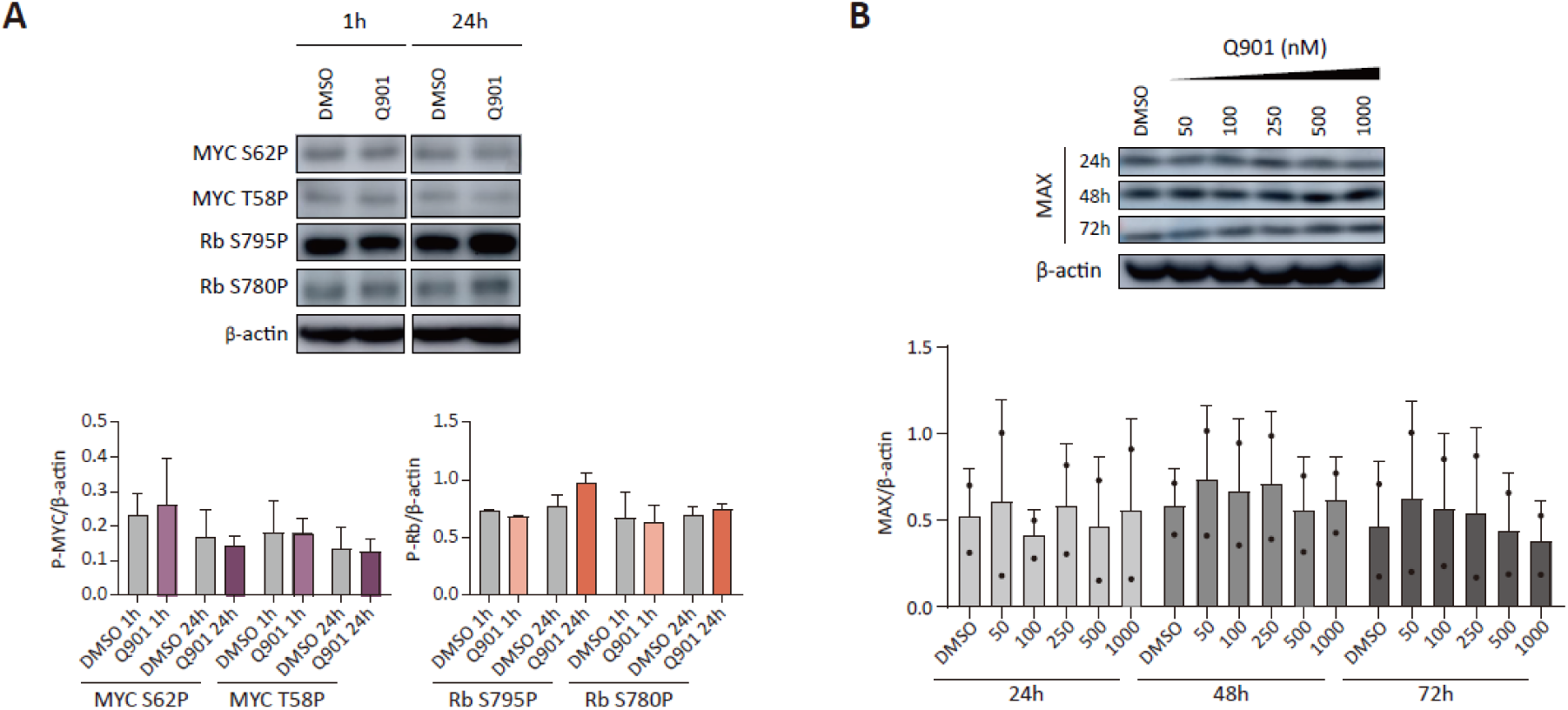
Expression levels of MYC and E2F1 activity-related genes. (**A**) Western blot data of phospho-MYC and phospho-Rb after Q901 treatment (n = 2 or 3; two-way ANOVA followed by Šidák’s multiple comparison test, data represent mean ± SD). (**B**) Western blot data of MAX expression after Q901 treatment (n = 2; two-way ANOVA followed by Šidák’s multiple comparison test, data represent mean ± SD).

**Fig. S6.**
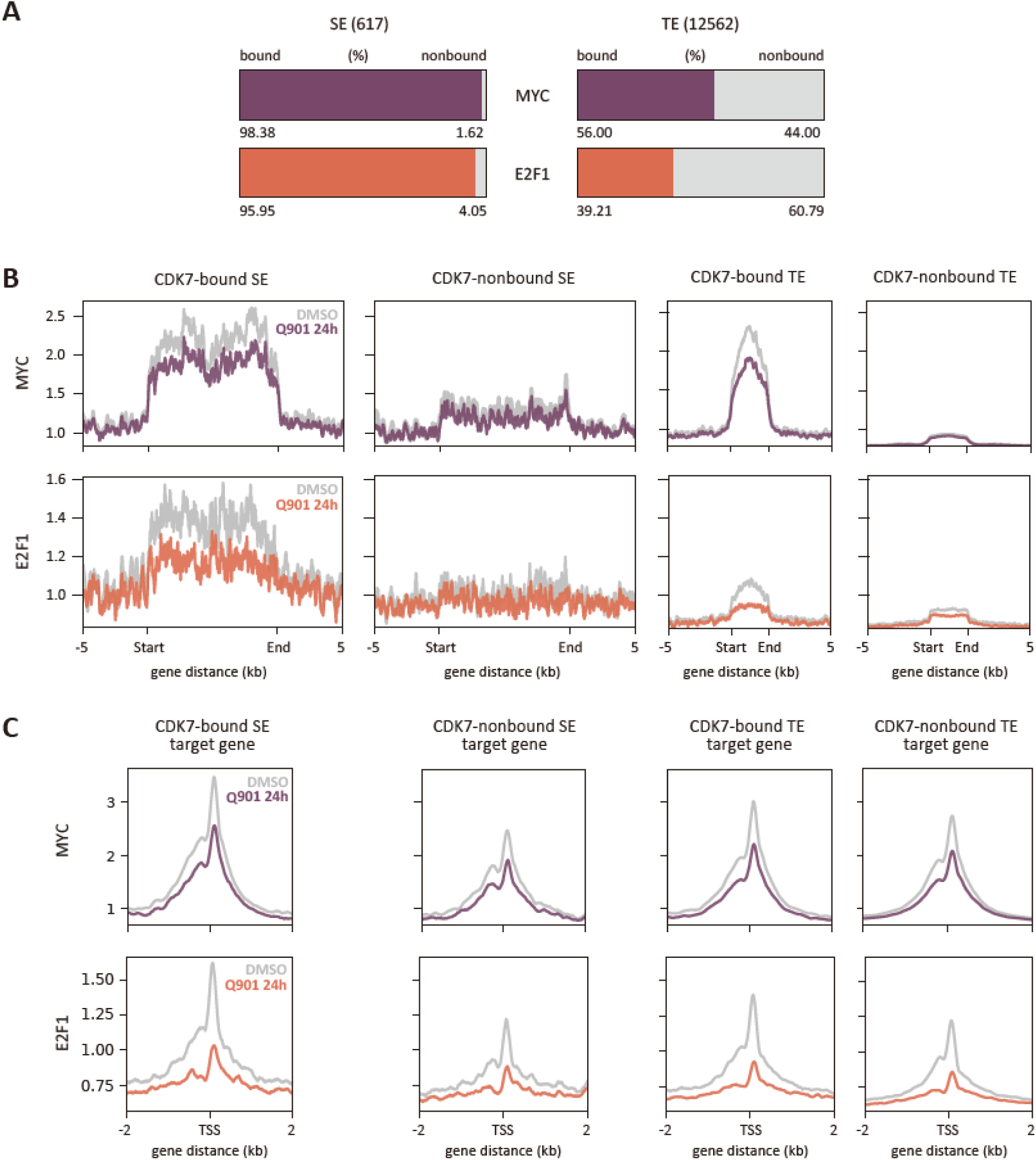
MYC and E2F1 binding profiles across enhancer regions and their target genes. (**A**) Box plot showing the ratio of MYC/E2F1-bound and non-bound regions in SEs, TEs. (**B**) Average plots of MYC and E2F1 ChIP-seq signals across the four enhancer groups. (**C**) Average plots of MYC and E2F1 ChIP-seq signals across the four groups of enhancer target protein-coding genes.

**Fig. S7.**
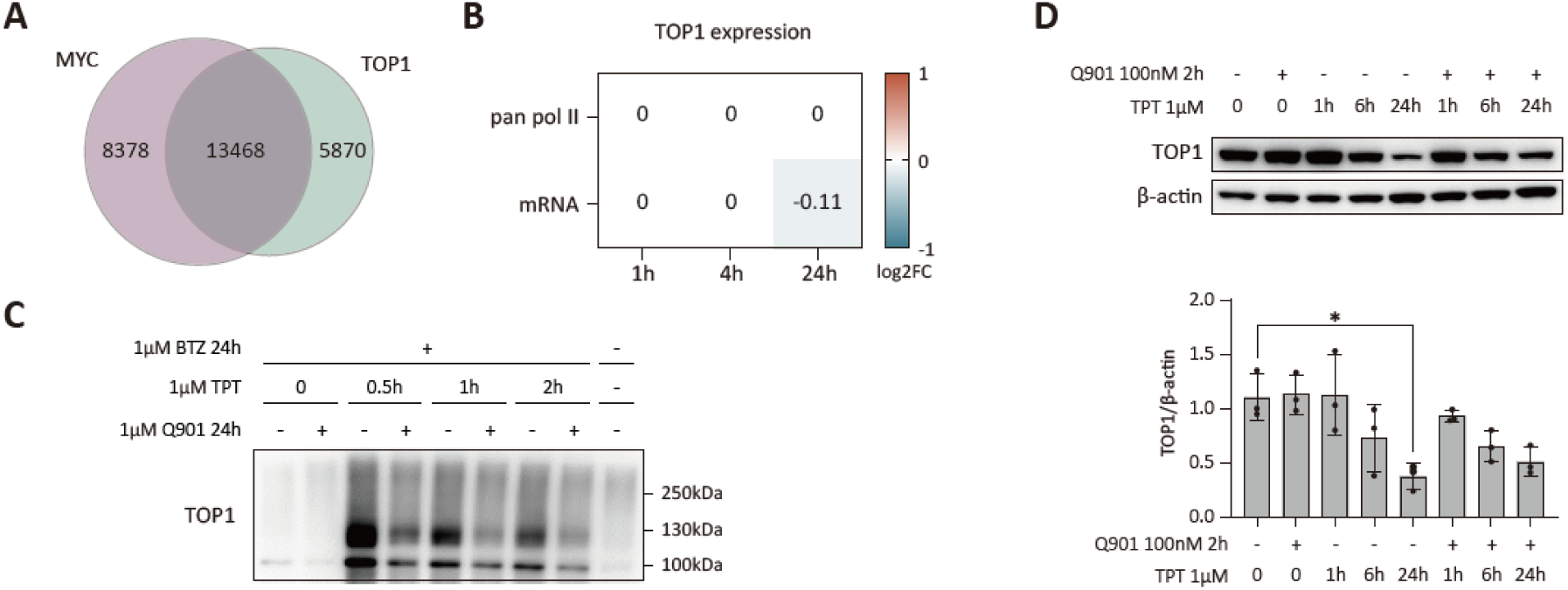
Transcriptional and protein-level regulation of TOP1 in response to Q901 and TPT treatments. (**A**) Venn diagram shows the overlap between MYC-bound and TOP1-bound genes. (**B**) Heatmap displays log2FC (Q901/DMSO) from pan RNAPII ChIP-seq and mRNA-seq data for TOP1. (**C**) Detection of TOP1-DPCs by DUST assay. (**D**) MCF-7 cells were pretreated with Q901 or DMSO for 2 h, followed by treatment with TPT for the indicated time points. Cell lysates were subjected to western blot assay using anti-TOP1 antibody (left). Bar graph (right) shows quantification of TOP1 normalized to β-actin (n = 3; two-way ANOVA followed by Šidák’s multiple comparison, data represent mean ± SD).

**Fig. S8.**
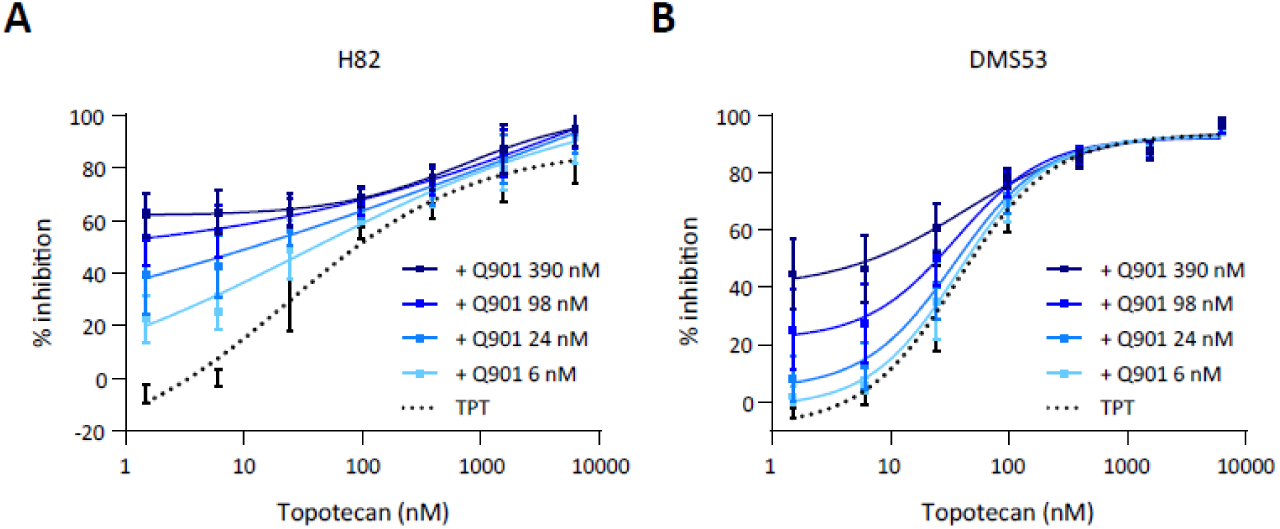
Synergistic effect of Q901 in combination with a TOP1 inhibitor. (**A** and **B**) Human SCLC cells, H82 (A), and DMS53 (B), were treated with varying concentrations of Q901 in combination with TPT for 72 h. Cell viability was measured using the CellTiter-Glo assay system. The percentage of cell growth inhibition was quantified by normalizing luminescence readings to untreated controls (n = 2, data represent mean ± SD).

**Fig. S9.**
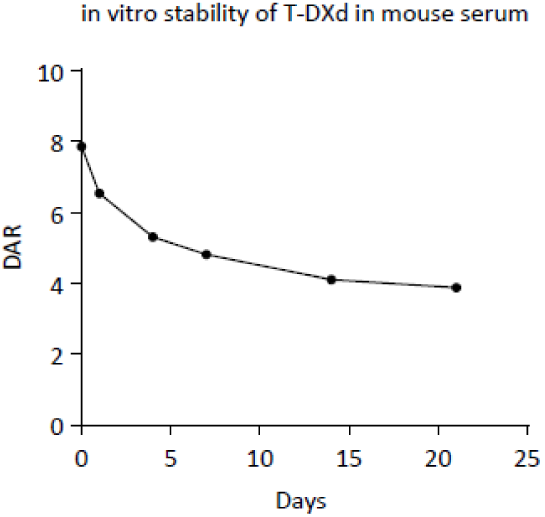
Serum stability and DAR profiling of T-DXd over time. Serum stability and DAR profiling of T-DXd over time. A gradual shift in the DAR of T-DXd from 8 to 4 was observed by day 21, indicating sustained payload release in mouse serum.

**Table S1.** The selectivity of Q901 against different CDK/Cyclin pairs. N/A: Not Applicable.

| CDKs | Residual activity at 1μM<br>(% of control) | IC<sub>50</sub>, (μM) |
| --- | --- | --- |
| CDK7/CycH/MAT1 | 1 | 0.01 |
| CDK1/CycA2 | 90 | 8.71 |
| CDK1/CycB1 | 93 | >10 |
| CDK1/CycE1 | 92 | 5.00 |
| CDK12/CycK | 84 | 2.48 |
| CDK13/CycK | N/A | 7.69 |
| CDK16/CycY | 95 | 6.94 |
| CDK19/CycC | 105 | >10 |
| CDK2/CycA2 | 91 | >10 |
| CDK2/CycE1 | 97 | 8.09 |
| CDK3/CycC | 85 | >10 |
| CDK3/CycE1 | 111 | >10 |
| CDK4/CycD1 | 96 | >10 |
| CDK4/CycD3 | 91 | >10 |
| CDK5/p25NCK | 93 | >10 |
| CDK5/p35NCK | 93 | >10 |
| CDK6/CycD1 | 96 | >10 |
| CDK6/CycD3 | 100 | >10 |
| CDK8/CycC | 86 | >10 |
| CDK9/CycK | 86 | 8.58 |
| CDK9/CycT1 | 91 | >10 |

