## supplemental figurea and table for "Sensitizing tumor response to topoisomerase I antibody drug conjugate by selective CDK7 inhibition"

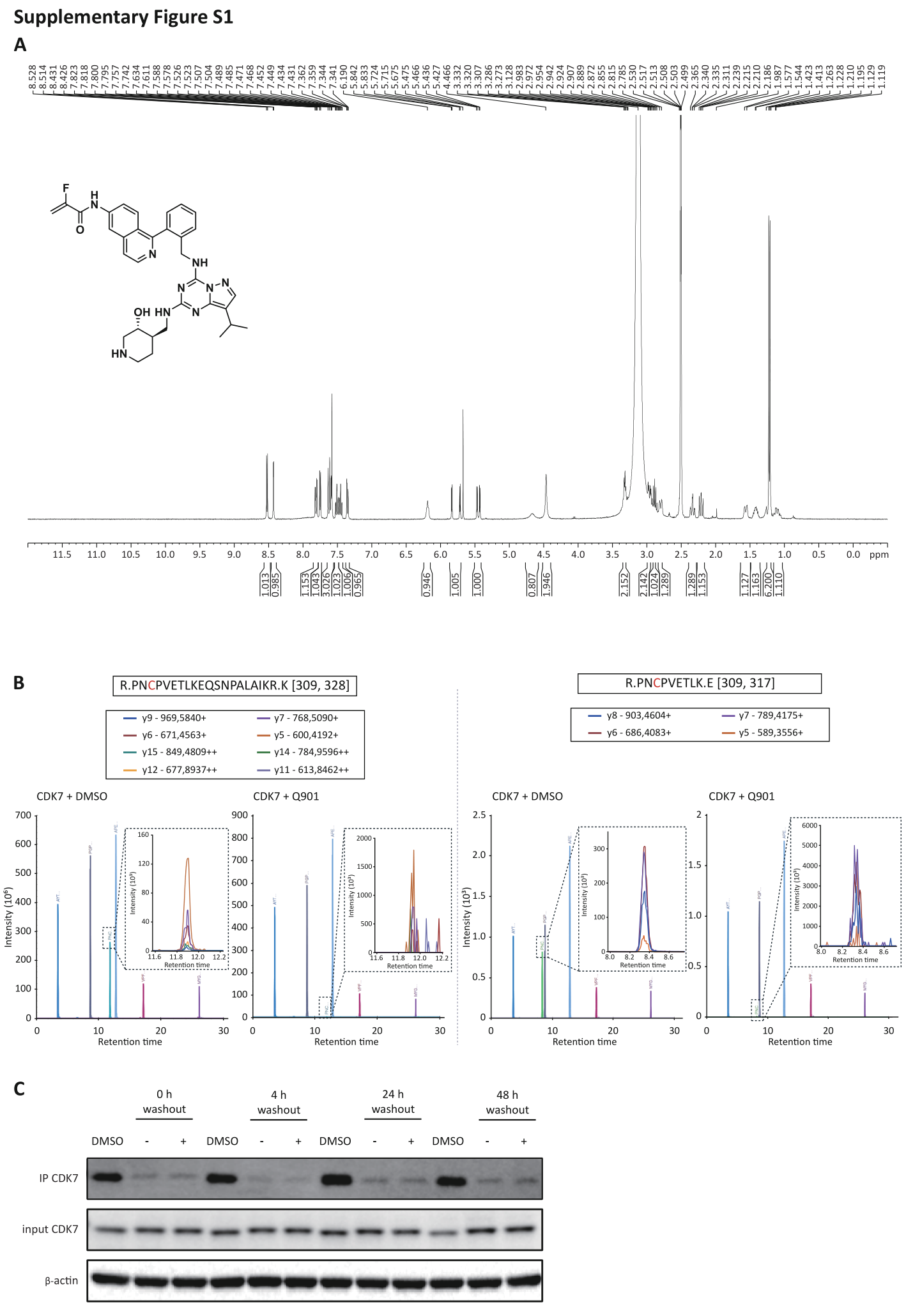

Fig. S1. Biochemical and proteomic validation of Q901 specificity.

(**A**) ^1^H NMR spectrum of Q901 was acquired using variable temperature (VT) NMR in DMSO-d₆. (**B**) Targeted proteomics analysis to determine the Q901 binding sites on CDK7. The recombinant CAK trimeric complex was incubated with Q901 or DMSO, followed by protease digestion and peptide mapping via LC-MS/MS. Chromatograms show peptide fragments generated by ArgC (Clostripain) digestion (right) and ArgC/Trypsin digestion (left). The expanded boxes highlight the peak of C312 containing peptides, which are reduced following Q901 treatment, indicating covalent modification at this site. (**C**) Representative Western blot images from the pulse-chase assay described in Fig. 1F. A2780 cells were treated with 6 nM Q901 for 4 h and then divided into two groups. One group (- wash out) remained in the Q901-containing medium for continuous incubation, while the other group (+ wash out) underwent a drug washout, where the medium was completely removed and replaced with fresh drug-free medium before further incubation for the indicated times. Bio-QS-labeled CDK7 was immunoprecipitated using streptavidin agarose beads (SA), and the levels of free CDK7 were analyzed by immunoblotting. These images in this figure were quantified in Fig. 1F.

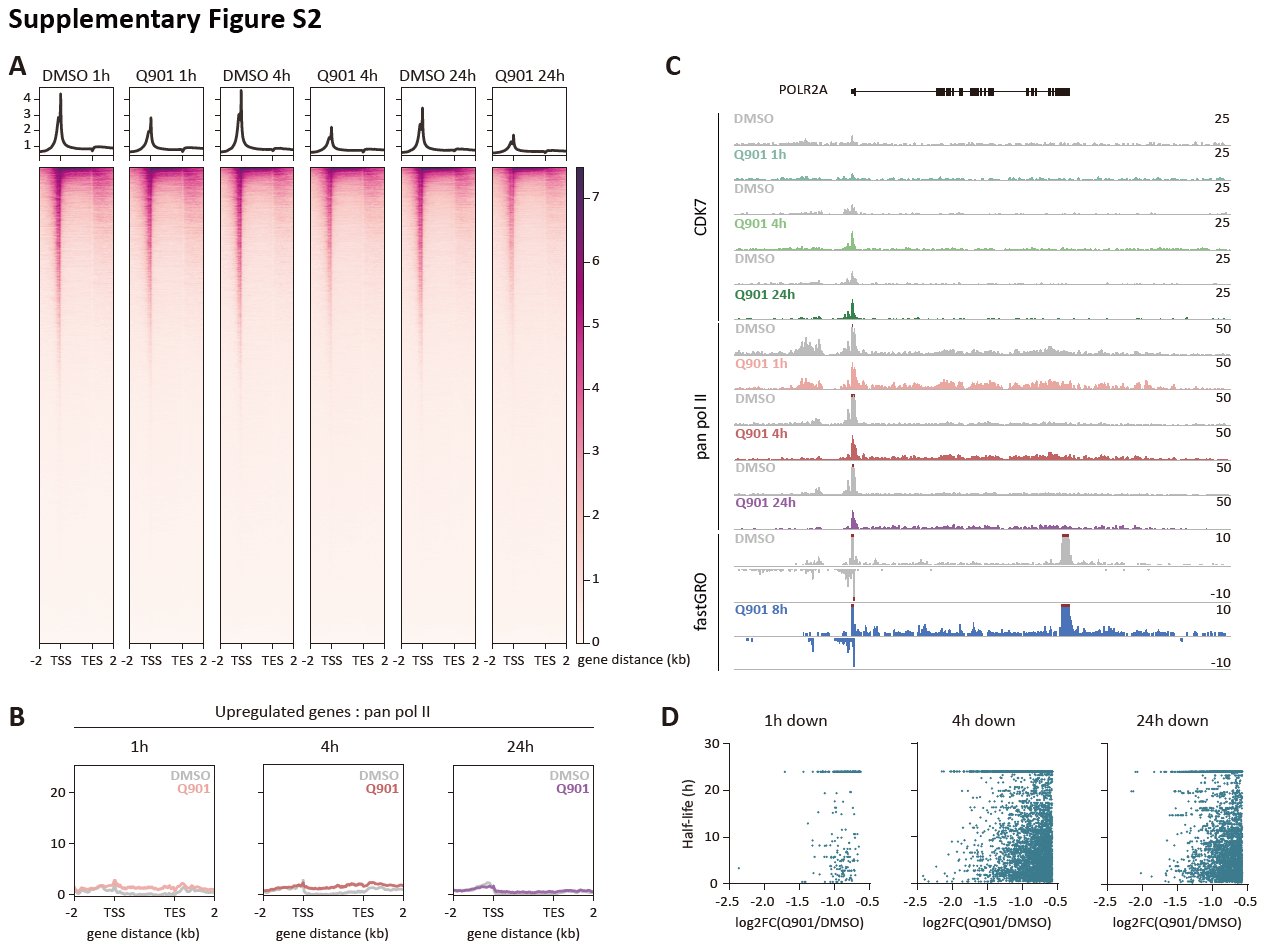

Fig. S2. Global and gene-specific changes in RNAPII occupancy following Q901 treatment.

(**A**) Heatmap of pan RNAPII ChIP-seq for all annotated genes (n = 2, p-value ≤ 0.05). (**B**) Average ChIP-seq plots of pan RNAPII for upregulated genes across various time points following Q901 treatment, presented on the same scale as those for downregulated genes in Fig. 2D. (**C**) Track image of POLR2A, an example of an upregulated gene by Q901 treatment. (**D**) Scatter plots showing the correlation between the mRNA half-lives of downregulated genes and log2FC values of pan RNAPII ChIP-seq for the corresponding genes.

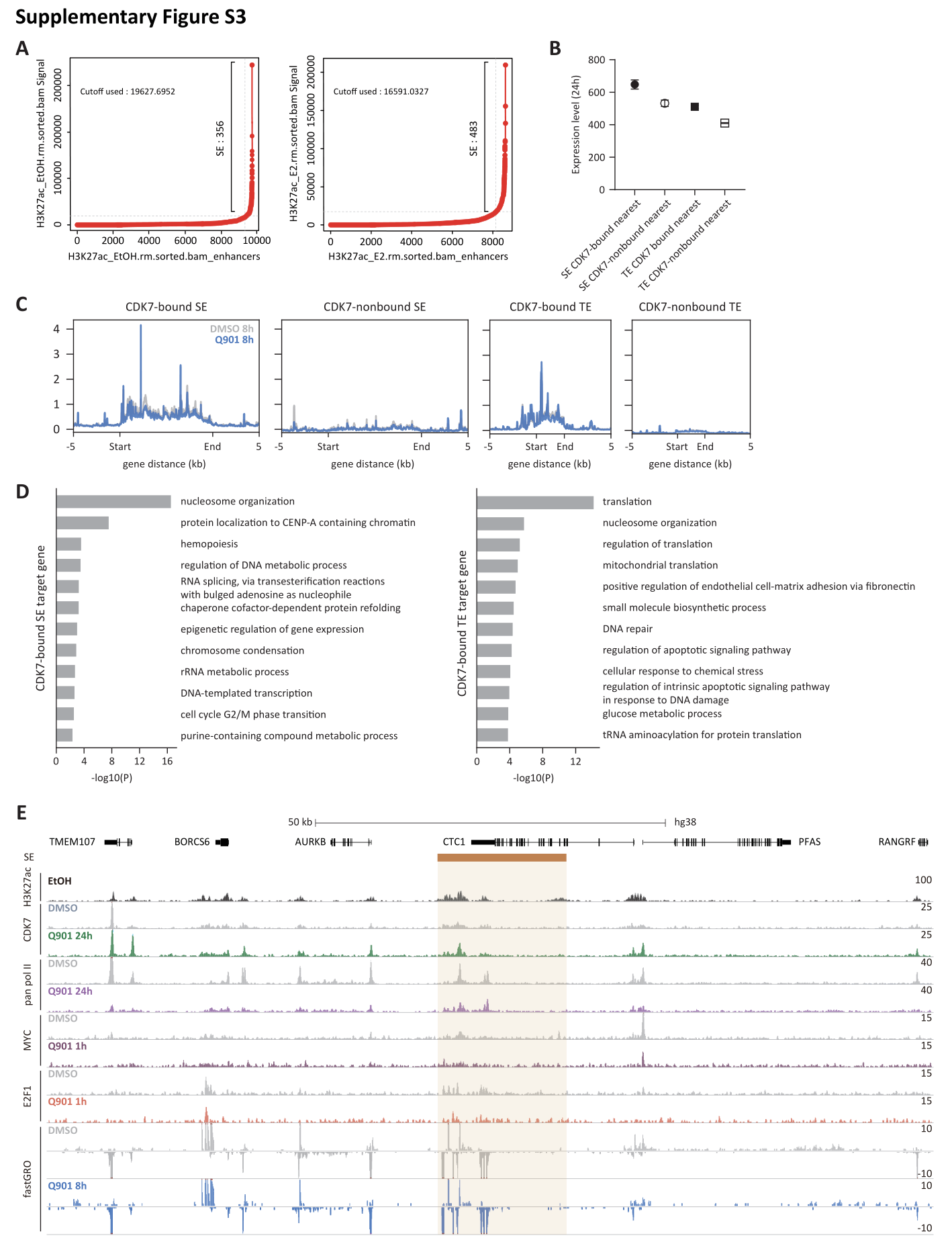

Fig. S3. Transcription profiles of enhancer target genes.

(**A**) The results of SE calling using the ROSE program with H3K27ac ChIP-seq (GSE62229). (**B**) Expression levels of enhancer target genes (pan RNAPII ChIP-seq; n = 2, Q901 1h treatment condition, data represent mean ± SEM). (**C**) Average fastGRO signals of four enhancer groups. (**D**) GO analysis results of target genes regulated by CDK7-bound SE and CDK7-bound TE. (**E**) Track images showing ChIP-seq signals for H3K27ac, CDK7, pan RNAPII, MYC, and E2F1, along with fastGRO, at a representative CDK7-bound SE region (highlighted in yellow) and it associated target genes.

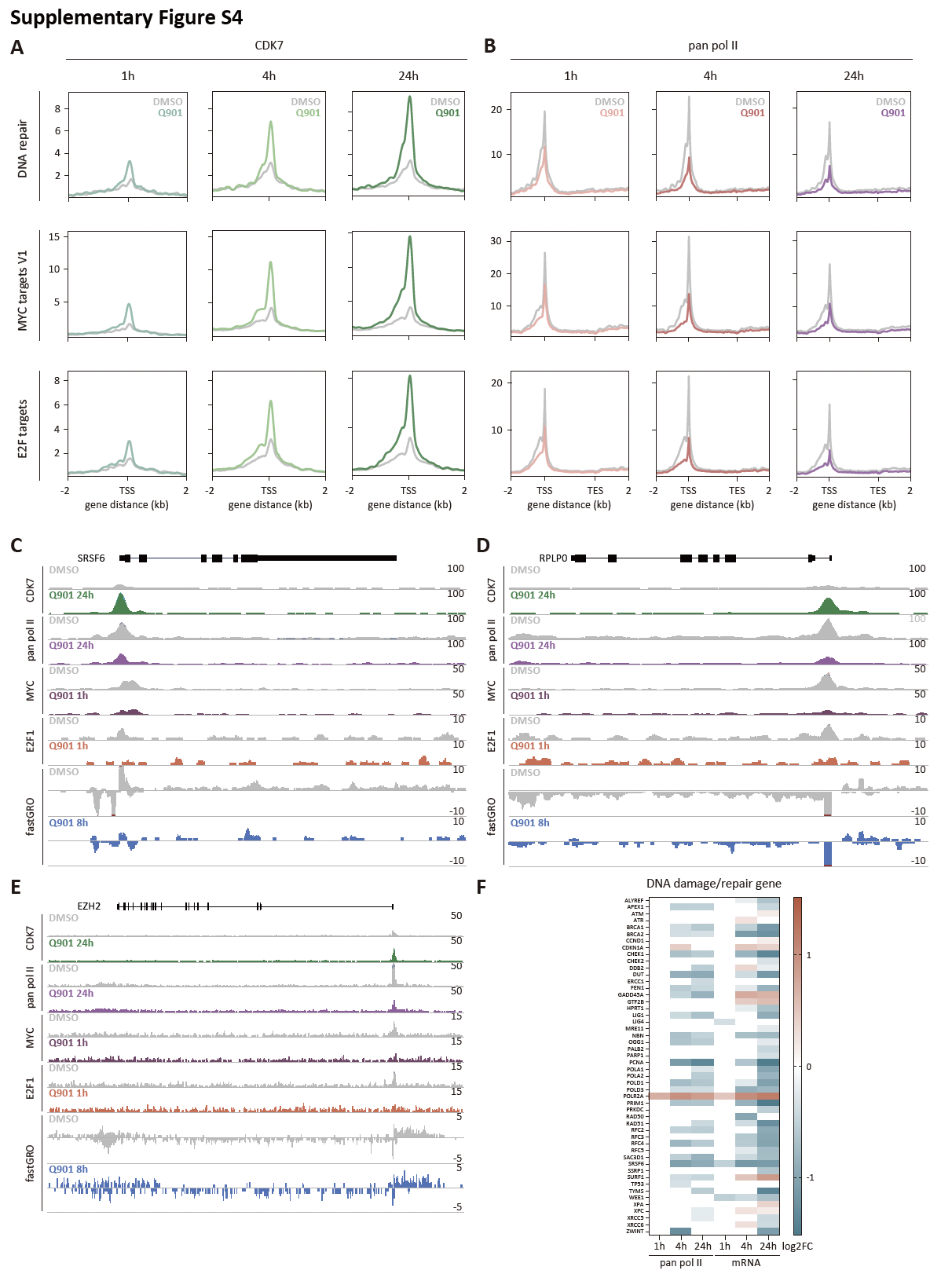

Fig. S4. CDK7 and RNAPII ChIP-seq signal at DNA Repair, MYC, and E2F target genes.

(**A** and **B**) Average ChIP-seq signal plots of CDK7 (A) and pan RNAPII (B) for gene sets related to DNA repair, MYC targets V1, and E2F targets. (**C**) Track image of SRSF6 gene, a representative gene from the DNA repair pathway. (**D**) Track image of RPLP0 gene, a representative gene from the MYC targets V1 pathway. (**E**) Track image of EZH2 gene, a representative gene from the E2F targets pathway. (**F**) Heatmap showing log2FC values of DNA damage/repair genes expression from pan RNAPII ChIP-seq and mRNA-seq data (right; n = 3, FDR ≤ 0.1).

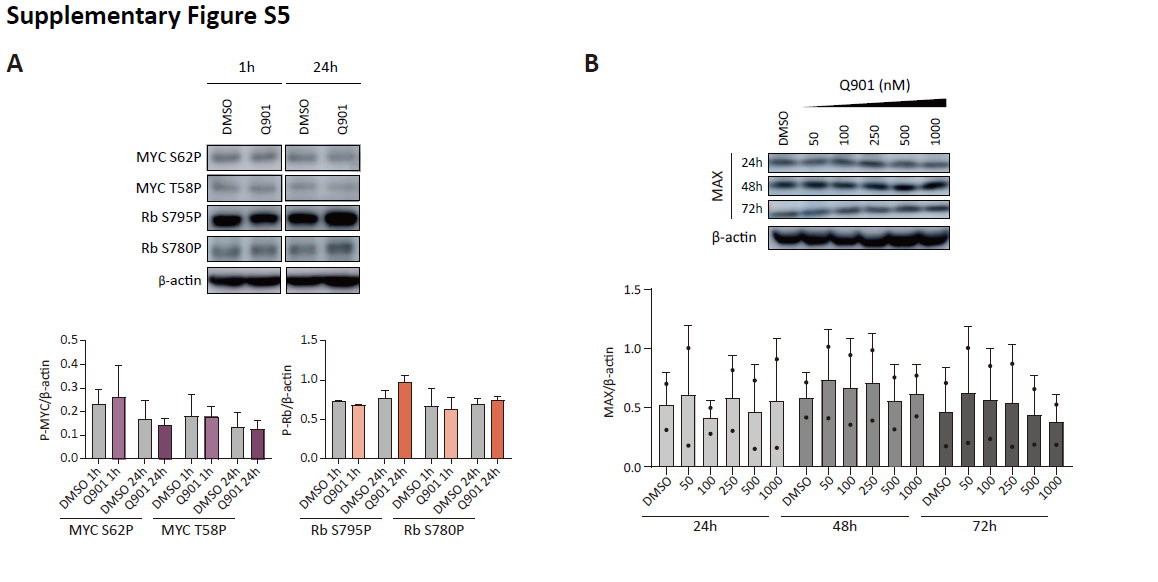

Fig. S5. Expression levels of MYC and E2F1 activity-related genes.

(**A**) Western blot data of phospho-MYC and phospho-Rb after Q901 treatment (n = 2 or 3; two-way ANOVA followed by Šidák's multiple comparison test, data represent mean ± SD). (**B**) Western blot data of MAX expression after Q901 treatment (n = 2; two-way ANOVA followed by Šidák's multiple comparison test, data represent mean ± SD).

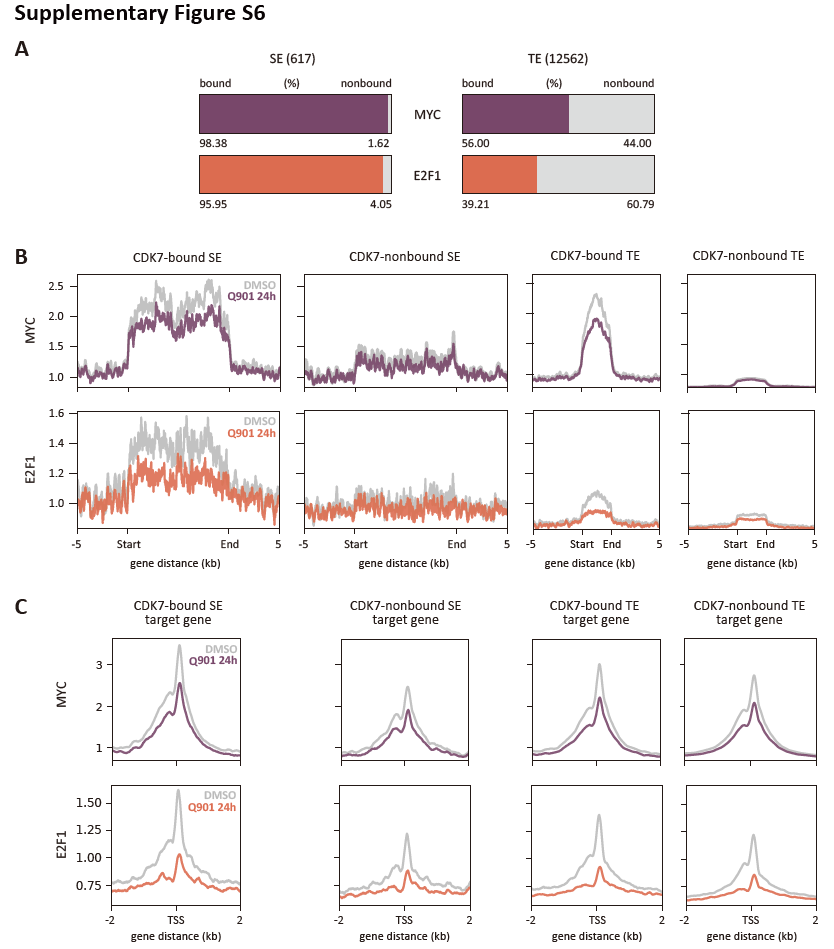

Fig. S6. MYC and E2F1 binding profiles across enhancer regions and their target genes.

(**A**) Box plot showing the ratio of MYC/E2F1-bound and non-bound regions in SEs, TEs. (**B**) Average plots of MYC and E2F1 ChIP-seq signals across the four enhancer groups. (**C**) Average plots of MYC and E2F1 ChIP-seq signals across the four groups of enhancer target protein-coding genes.

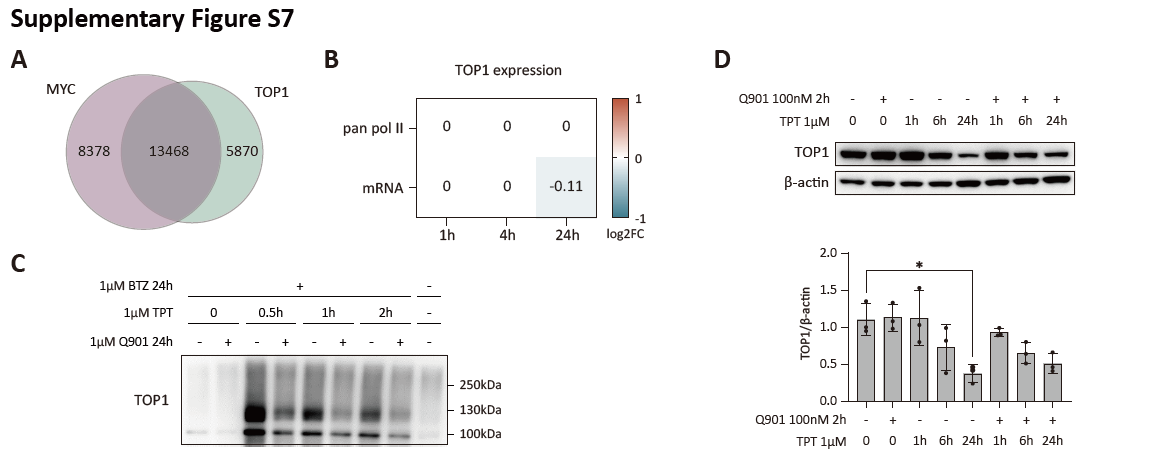

Fig. S7. Transcriptional and protein-level regulation of TOP1 in response to Q901 and TPT treatments.

(**A**) Venn diagram shows the overlap between MYC-bound and TOP1-bound genes. (**B**) Heatmap displays log2FC (Q901/DMSO) from pan RNAPII ChIP-seq and mRNA-seq data for TOP1. (**C**) Detection of TOP1-DPCs by DUST assay. (**D**) MCF-7 cells were pretreated with Q901 or DMSO for 2 h, followed by treatment with TPT for the indicated time points. Cell lysates were subjected to western blot assay using anti-TOP1 antibody (left). Bar graph (right) shows quantification of TOP1 normalized to β-actin (n = 3; two-way ANOVA followed by Šidák's multiple comparison, data represent mean ± SD).

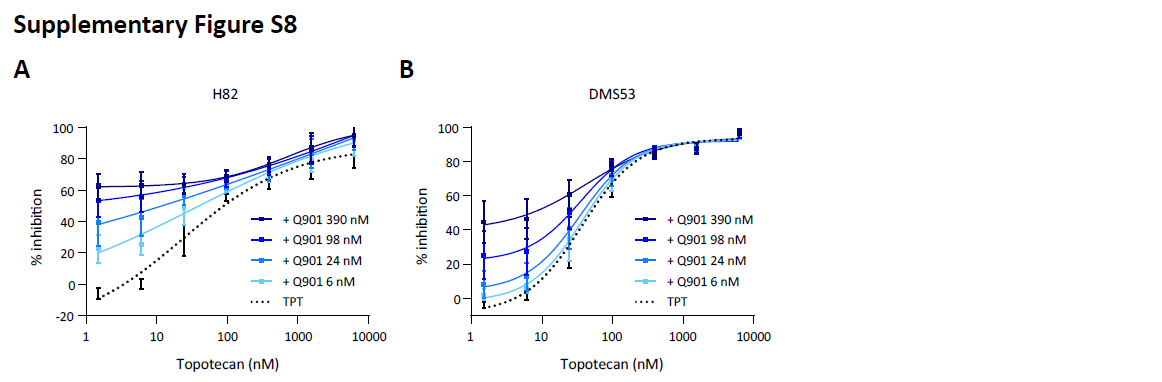

Fig. S8. Synergistic effect of Q901 in combination with a TOP1 inhibitor.

(**A** and **B**) Human SCLC cells, H82 (A), and DMS53 (B), were treated with varying concentrations of Q901 in combination with TPT for 72 h. Cell viability was measured using the CellTiter-Glo assay system. The percentage of cell growth inhibition was quantified by normalizing luminescence readings to untreated controls (n = 2, data represent mean ± SD).

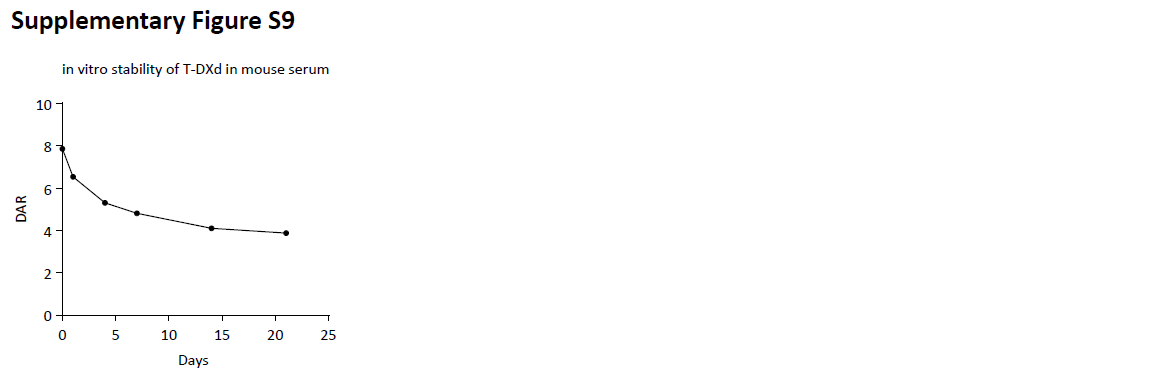

Fig. S9. Serum stability and DAR profiling of T-DXd over time. Serum stability and DAR profiling of T-DXd over time.

A gradual shift in the DAR of T-DXd from 8 to 4 was observed by day 21, indicating sustained payload release in mouse serum.

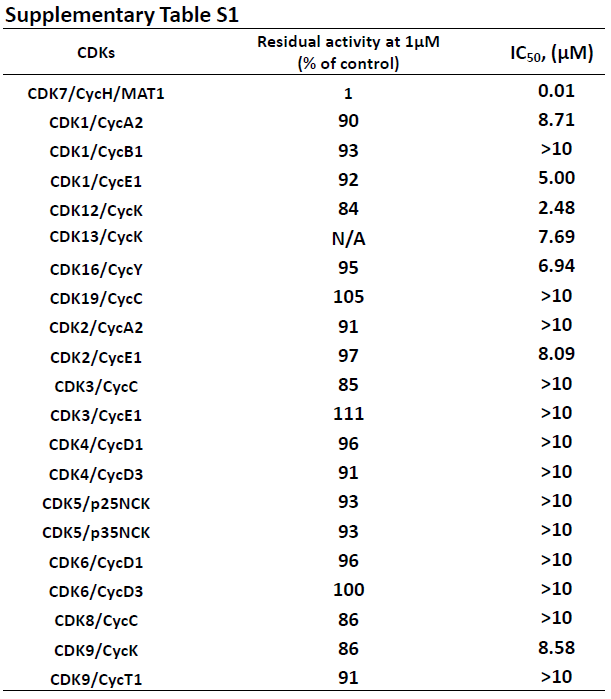

Table S1. The selectivity of Q901 against different CDK/Cyclin pairs.

N/A: Not Applicable.
